# Beta-coalescents when sample size is large

**DOI:** 10.64898/2025.12.30.697022

**Authors:** Jonathan A. Chetwynd-Diggle, Bjarki Eldon

## Abstract

Individual recruitment success, or the offspring number distribution of a given population, is a fundamental element in ecology and evolution. Sweepstakes reproduction refers to a highly skewed individual recruitment success without involving natural selection and may apply to individuals in broadcast spawning populations characterised by Type III survivorship. We consider an extension of the Schweinsberg (2003) model of sweepstakes reproduction for a haploid panmictic population of constant size *N*; the extension also works as an alternative to the Wright-Fisher model. Our model incorporates an upper bound on the random number of potential offspring (juveniles) produced by a given individual. Depending on how the bound behaves relative to the total population size, we obtain the Kingman (1982a,c,b) coalescent, an incomplete Beta-coalescent, or the (complete) Beta-coalescent of Schweinsberg (2003). We argue that applying such an upper bound is biologically reasonable. Moreover, we estimate the error of the coalescent approximation. The error estimates reveal that convergence can be slow, and small sample size can be sufficient to invalidate convergence, for example if the stated bound is of the form *N/* log *N*. We use simulations to investigate the effect of increasing sample size on the site-frequency spectrum. When the limit is a Beta-coalescent, the site frequency spectrum will be as predicted by the limiting tree even though the full coalescent tree may deviate from the limiting one. When in the domain of attraction of the Kingman coalescent the effect of increasing sample size depends on the effective population size as has been noted in the case of the Wright-Fisher model. Conditioning on the population ancestry (the random ancestral relations of the entire population at all times) may have little effect on the site-frequency spectrum for the models considered here (as evidenced by simulation results).

## 1 Introduction

A key element in much of mathematical population genetics is recruitment success, or the distribution of the number of offspring (gene copies), produced by each individual gene copy at any given time. From models of offspring number distribution, or individual recruitment success, one develops appropriate statistical tools for inferring evolutionary histories of natural populations. Commonly, data takes the form of the DNA sequence of a given genetic locus (seen here as a contiguous non-recombining segment of a chromosome) for each individual in a sample from the population. We use differences between these sequences to infer information about the genealogical tree relating the sequences in the sample. If we know the distribution of the genealogical trees under our population models, we can aim to use DNA sequence data to distinguish between evolutionary scenarios.

The celebrated Kingman coalescent (Kingman, 1982a,c,b) provides a reference point. It arises as an approximation to the (unknown) genealogical tree relating ancestors of a population genetic sample drawn from a natural population, and when the approximate trees are viewed on an appropriate timescale. The timescale is determined by the ‘effective population size’, denoted *N*_*e*_. The Kingman coalescent is a good approximation to the genealogy of a sample for a whole raft of population models, primarily characterised by ‘small’ family sizes (discussed more precisely below), and it is this stability under small perturbations of the underlying population model that makes it a useful tool for inference.

A key feature of Kingman’s coalescent is that at most two ancestral lineages merge in a given coalescence event. This reflects two underlying assumptions: first, the sample size is a negligible fraction of *N*_*e*_; second, in the corresponding population models, any given individual has a negligibly small number of offspring relative to *N*_*e*_. Regarding the first assumption, in many cases *N*_*e*_ has been estimated to be much smaller than census population size (Árnason, 2004; Tenesa et al., 2007). With progress in DNA sequencing technology, sample sizes for population genetic studies are growing rapidly (Bycroft et al., 2018), so that the stated assumption about sample size can increasingly be called into question. In addition, advances in algorithm design enable efficient simulations of genealogical trees of large samples (Kelleher et al., 2016). Regarding the second assumption, observations from ecology and empirical population genetics suggest that in highly fecund species (ranging from fungi to gadids) family sizes may be highly variable in any given generation. At any given time a small number of individuals may produce numbers of *potential offspring* (offspring that may survive to reproducing age) proportional to the population size (May, 1967; Hedgecock, 1994a; Li and Hedgecock, 1998; Hedgecock and Pudovkin, 2011). These organisms are characterised by broadcast spawning, external fertilisation, extremely high fecundity and a so-called Type III survivorship curve (high initial mortality followed by a slow decrease in survival probability). To counter the high initial mortality populations may evolve according to ‘sweepstakes reproduction’, which is a mechanism turning high fecundity into a skewed individual recruitment success. In sweepstakes a few individuals occasionally match by chance reproduction with favorable environmental conditions and thus contribute a substantial fraction of the surviving offspring at such a time (Beckenbach, 1994; Hedgecock, 1994a; Hedgecock et al., 2007; Li and Hedgecock, 1998; Hedgecock and Pudovkin, 2011; Williams, 1975; Árnason et al., 2023). This kind of recruitment dynamics has also been referred to as ‘random sweepstakes’ (Árnason et al., 2023) to distinguish from a different mechanism involving natural selection for generating skewed recruitment success (Williams, 1975). Sweepstakes reproduction has been invoked to help explain chaotic genetic patchiness observed in marine populations (Iannucci et al., 2020; Vendrami et al., 2021), and lower observed genetic diversity than expected based on estimates of census population size (Hedgecock, 1994b; Hedgecock and Pudovkin, 2011). Thus, sweepstakes may play a substantial role in connectivity among marine populations, which in turn is fundamental to population, metapopulation, and community dynamics and structure, genetic diversity, and resilience to anthropogenic impact (Selkoe et al., 2016; Gagnaire et al., 2015; Cowen and Sponaugle, 2009; Cowen et al., 2006; Botsford et al., 2001).

Examples of populations potentially characterised by sweepstakes reproduction include Atlantic cod (*Gadus morhua*), Pacific oyster (*Crassostrea gigas*), red drum (*Sciaenops ocellatus*), Japanese sardines, hydrothermal vent taxa, and the Antarctic limpet (*Nacella concinna*) (Árnason, 2004; Árnason and Halldórsdóttir, 2015; Niwa et al., 2016; Turner et al., 2002; Li and Hedgecock, 1998; Hurtado et al., 2004; Vendrami et al., 2021; Árnason et al., 2023). Similar considerations apply to, for example, plant populations which distribute pollen, and insect populations where individuals of one gender far outnumber those of the other. In populations characterised by sweepstakes reproduction type frequencies are in the domain of attraction of forward-in-time Fleming-Viot measure-valued (Fleming and Viot, 1979; Ethier and Kurtz, 1993) processes admitting jumps in the type frequency (Birkner and Blath, 2009; Huillet and Möhle, 2011; Möhle, 2001; Donnelly and Kurtz, 1999; Bertoin and Le Gall, 2003, 2005, 2006), and sample gene genealogies are in the domain of attraction of *multiple merger* coalescents where a random number of ancestral lineages (ancestors of sampled gene copies) merges whenever mergers occur (Sagitov, 1999; Pitman, 1999; Donnelly and Kurtz, 1999; Schweinsberg, 2000; Möhle and Sagitov, 2001; Schweinsberg, 2003; Sagitov, 2003). Multiple-merger coalescents are referred to as Λ-coalescents (Lambda-coalescents) if at most one group of a random number of ancestral lineages merges at a time (asynchronous mergers), and Ξ-coalescents (Xi-coalescents) if at least two such mergers can occur at the same time (simultaneous mergers). It has been suggested that multiple-merger coalescents provide a more appropriate description, compared to the Kingman (1982a,c,b) coalescent, of the genealogy of a sample from a highly fecund population characterised by sweepstakes reproduction (Eldon and Wakeley, 2006; Árnason, 2004; Sargsyan and Wakeley, 2008; Birkner et al., 2011; Árnason and Halldórsdóttir, 2015; Birkner and Blath, 2008; Birkner et al., 2013; Árnason et al., 2023). A subfamily of Λ-coalescents, known as Beta-coalescents, has been investigated to some extent (Schweinsberg, 2003; Birkner et al., 2005; Dhersin and Yuan, 2015; DAHMER et al., 2014; Berestycki et al., 2008a, 2007). Beta-coalescents are parametrised by a quantity *α >* 0, which captures the right-tail behaviour (see Equation (1.1)) of the distribution of the random number of potential offspring (see Definition 2.3) contributed by a single individual in a population model of sweepstakes reproduction (Schweinsberg, 2003). For *α* ∈ (1, 2) (we exclude the case 0 *< α* ≤ 1), the Beta-coalescent arises naturally from a haploid population model that we shall refer to as the *Schweinsberg model* (Schweinsberg, 2003). The Schweinsberg model is a model of discrete (non-overlapping) generations, in which all current individuals have a chance of producing offspring in any given generation. Thus, the Schweinsberg model shares certain features with the Wright-Fisher model (Wright, 1931; Fisher, 1922). However, a key deviation from the Wright-Fisher model is the production of potential offspring according to a non-trivial law; a given number of the potential offspring is then sampled without replacement (conditional on there being enough of them) to survive to maturity. It is assumed that, with *X* denoting the random number of juveniles produced by a given individual (*α, C >* 0 fixed),

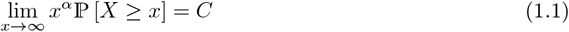

(Schweinsberg, 2003, Equation 11). The value of *α* in (1.1) determines the timescale of coalescence, i.e. how to calibrate time so that the limiting coalescent becomes a good description of the (random) gene genealogy of a sample from a large population evolving according to a given model. Denote by *c*_*N*_ the probability that two individuals, sampled at random and at the same time from the population, have a common parent in the previous generation. Then as *N→∞* and *α* ∈ (1, 2), *N* ^*α*−1^*c*_*N*_ *→ αCB*(2 −*α, α*)*/*(𝔼 [*X*])^*α*^, where *B*(2 −*α, α*) is the beta function (Schweinsberg, 2003). Measuring time in units of 1*/c*_*N*_ generations, as *N→∞*, the genealogy of a finite sample from the population converges, in the sense of finite dimensional distributions, to a Beta coalescent with parameter *α* (Schweinsberg, 2003).

The key assumption in (1.1) is that the number of potential offspring produced by any given individual can be arbitrarily large. An obvious objection to the Schweinsberg model as a justification of the Beta-coalescent model is that *α* in (1.1) is determined by the behaviour of the distribution of unrealistically high numbers of potential offspring. Even though some organisms can produce vast numbers of such offspring, they can’t produce arbitrarily many. The importance of this restriction may become clearer when considering diploid populations (e.g. Birkner et al. (2018); Möhle and Sagitov (2003)), where the offspring represent (at least) fertilised eggs. Thus, it is plausible that at least for some diploid populations (e.g. broadcast spawning marine organisms), the number of fertilised eggs from a given parent pair can be at most a fraction of the population size. As we will see, incorporating this restriction makes a difference to predictions about genetic diversity.

We will say that a coalescent approximation ‘breaks down’ when a given coalescent stops being a good approximation to the exact genealogy of a sample drawn from a finite population; we will discuss this in the context of increasing sample size and sweepstakes, and identify the sample size at which a given multiple-merger coalescent approximation breaks down. The Kingman coalescent breaks down as an approximation to the genealogy of a sample from a population of finite size and small family sizes when the sample size is large enough (see e.g. Bhaskar et al. (2014); Wakeley and Takahashi (2003); Fu (2006); Melfi and Viswanath (2018a,b)). Similarly, one might expect gene genealogies of large enough samples from a finite population evolving under the Schweinsberg model to deviate from predictions of Beta-coalescents.

Our aims in this paper are threefold. First we shall consider an extension of the Schweinsberg model, and incorporate an upper bound on the random number of potential offspring that can be produced by a single individual. The bound will depend on the total population size. We then identify the corresponding coalescents. Depending on the bound, this will be a *(i)* complete Beta coalescent, *(ii)* an *incomplete* Beta coalescent, or *(iii)* the Kingman coalescent. Second, we shall investigate the variability in *c*_*N*_ for large *N*. This provides some insight into how variable estimates of (coalescent) effective population size (Definition 2.5) will be for a population evolving under this model. Third, we will use informal arguments and simulations to investigate the breakdown in the coalescent approximation for large sample sizes. In this context, what we really care about is when the breakdown will be evident in data. We shall argue that, in practice, this will occur at a larger sample size than that at which anomalies first appear in the full coalescent tree, but nonetheless at sample sizes not out of reach with current DNA sequencing technology.

The layout of the paper is as follows. In Section 2, we provide a background on Beta-coalescents (Section 2.1), the site frequency spectrum (Section 2.2), the Schweinsberg model (Definition 2.3), and previous studies on large sample size (Section 2.3). In Section 3 we introduce our extension of the Schweinsberg model (see (3.2) in Section 3.1), and give a precise statement of our main results, i.e. the rescaling of time required for convergence (as the population size *N→∞*) to a coalescent (i.e. the unit of time used to measure the time between events involving ancestors of the sample) (Proposition 3.2), and convergence to the coalescents (Propositions 3.3 and 3.4). In Sections 3.3 and 3.4 we consider the impact of increasing sample size; in Section 3.3 in connection to the Kingman coalescent; and in Section 3.4 we give arguments for the sample size required to see its effect on the site-frequency spectrum in the case of the Beta-coalescent of Schweinsberg (2003).

Section 4 is devoted to numerical experiments, and Section 5 summarizes and briefly discusses our main findings. Key lemmas are reviewed in Section 6, followed by proofs in Sections 7 to 10. Many of the ideas follow Schweinsberg (2003), although we have taken care to keep track of the error bounds, so that we can identify the sample size at which the coalescent approximation ‘breaks down’, i.e. when we can expect to start seeing deviations from the predicted genealogies. In particular, in Section 7 we give a proof of the timescaling for our model, in Section 9 we give a proof of convergence to the Kingman coalescent from our model, in Section 10 we give proofs of convergence to Beta-coalescents. In Appendices E and F we briefly describe algorithms used for the numerical examples in Section 4 and in Appendix D; Appendix C contains examples of predictions of Beta coalescents. Appendices A and B contain further discussions about the sample size needed to disrupt convergence to the Kingman coalescent. For background on population genetics and coalescent theory see for example Gale (1980), Gillespie (2004), Wakeley (2009), Berestycki (2009), Etheridge (2011), Ewens (2004), Durrett (2008)

## 2 Background

For ease of reference we collect together notation that will be used throughout.

### Definition 2.1

(Notation). *Let N* ∈ ℕ := {1, 2, … }*denote the population size. Asymptotics will be understood to hold as N→∞, unless otherwise noted*.

*Write* [*n*] := {1, 2, …, *n*} *for any natural number n* ∈ ℕ; *unless stated otherwise the sample size will be represented by n*.

*Given two positive sequences* (*x*_*i*_)_*i*∈ℕ_ *and* (*y*_*i*_)_*i*∈ℕ_ *with* (*y*_*i*_) *bounded away from zero write*

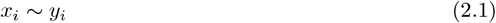

*if x*_*i*_*/y*_*i*_ → 1 *as i* → ∞. *When x*_*i*_*/y*_*i*_ → *c as i* → ∞ *where* 0 *< c <* ∞ *is fixed we write*

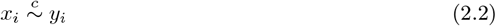

*In* (2.2) *the constant c is not specified and will change depending on what* (*x*_*i*_) *and* (*y*_*i*_) *are each time; when we use this notation we are empasizing the conditions under which* (2.2) *holds with* (*x*_*i*_) *and* (*y*_*i*_) *as given. For positive functions f, g such that* lim sup_*x*→∞_ *f* (*x*)*/g*(*x*) *<* ∞ *we write f* = 𝒪 (*g*).

*For any real number a and i* ∈ ℕ_0_ := {0, 1, 2, …} *we will write* (*a*)_0_ := 1 *and*

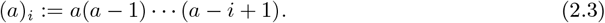

*For a given condition/event E we define*

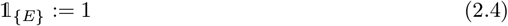

*if E holds, and take* 𝟙 _{*E*}_ = 0 *otherwise*.

*Let* 0 *< p* ≤ 1 *and a, b >* 0 *be fixed. We will write*

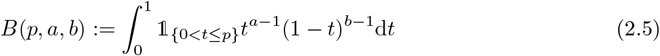

*It then holds that* 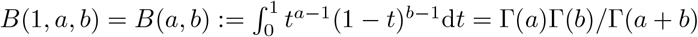.

### 2.1 Beta-coalescents

An ancestral process {*ξ*^*n,N*^} *≡* {*ξ*^*n,N*^ (*t*); *t ≥* 0} for a sample of *n* gene copies is a partition-valued Markov sequence on the partitions of [*n*] for any *n* ∈ ℕ describing the random ancestral relations of the sampled gene copies (we take a gene copy to be a contiguous non-recombining segment of a chromosome). The sample is assumed to be from a finite (haploid) panmictic population of size *N* evolving according to a given population model. One then aims to prove convergence (in the sense of finite-dimensional distributions) of {*ξ*^*n,N*^ (⌊*t/c*_*N*_ ⌋); *t ≥* 0 }to a continuous-time coalescent (provided *c*_*N*_ *→*0).

Beta-coalescents (Schweinsberg, 2003) and the coalescents considered by e.g. Eldon and Wakeley (2006) and Matuszewski et al. (2017) are special cases of Λ-coalescents, which were constructed independently by Donnelly and Kurtz (1999), Pitman (1999) and by Sagitov (1999), and which have since been studied by many authors (see Berestycki (2009) and Gnedin et al. (2014) for surveys). Like the Kingman coalescent, they are Markov processes, taking their values among partitions of ℕ. If {*ξ*(*t*), *t ≥*0} is a Λ-coalescent, we write {*ξ*^*n*^} ≡ {*ξ*^*n*^(*t*), *t ≥*0} for its restriction (itself Markov) to [*n*] for each *n* ∈ ℕ. The elements of [*n*] are used to arbitrarily label [*n*] individuals sampled from the population. Each block of the partition *ξ*^*n*^(*t*) (for any *t ≥* 0) corresponds to an individual ancestral to a subset of the sample of size *n* at time *t* before the present, and the elements of the block are the labels of the descendants of that ancestor in the sample. For example, *ξ*^*n*^(0) = {{1}, …, *n*}, and the partition {[*n*]} contains only the block [*n*] corresponding to an ancestor of the whole sample. In what follows, we shall abuse terminology and refer to the pure death process that records the total number of lineages ancestral to the sample at each time *t* before the present as the coalescent. If there are currently *k* ∈ {2, …, *n*} blocks in *ξ*^*n*^(*t*) then each transition involving *j* ∈ {2, …, *k*} of the blocks merging into one happens at rate *b*_*k,j*_ (which is *independent* of *n*), the remaining blocks are unchanged, and these are the only possible transitions. The rate of such mergers is not arbitrary; Pitman (1999) shows a one-to-one correspondence between these coalescents and finite measures on (0, 1]. For a given finite measure Λ_+_ on (0, 1], in the corresponding coalescent, suppose there are currently *k* lineages ancestral to a given sample. Then the rate at which a given subset of *j* of them coalesces into a single lineage is, with *a ≥* 0 fixed, and recalling (2.4) in Definition 2.1,

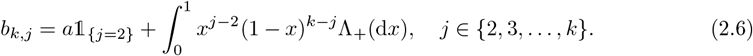

The Kingman coalescent is a Λ-coalescent with *a* = 1 and Λ_+_ = 0 in (2.6). Before we proceed we formally define the (complete) Beta-coalescent.

#### Definition 2.2

(The (complete) Beta coalescent (Schweinsberg, 2003)). *The Beta-coalescent with parameter α is a* Λ*-coalescent in which, if there are currently k lineages ancestral to the sample, the rate at which a particular subset of j of them coalesces into a single lineage is obtained from* (2.6) *by setting a* = 0 *and taking (recall* (2.5))

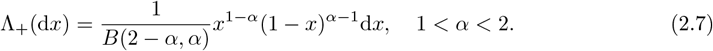

Even though one can take any strictly positive *α <* 2 in (2.7), the Beta-coalescent with *α <* 1 does not correspond to any (currently) known population model. In what follows, we shall restrict our attention to haploid populations. To incorporate diploidy would require Ξ-coalescents, which allow for simultaneous multiple mergers, with rates characterised through a measure Ξ on the infinite simplex

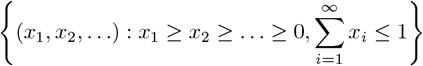

(Schweinsberg, 2000; Möhle and Sagitov, 2003; Sagitov, 2003; Birkner et al., 2013a; Blath et al., 2016; Birkner et al., 2018). Beta-coalescents describe the random ancestral trees of samples drawn from a population evolving according to the Schweinsberg (2003) model in which the law of the random family sizes has a specific tail behaviour.

#### Definition 2.3

(The Schweinsberg model (Schweinsberg, 2003)). *Consider a haploid panmictic population of fixed size N which evolves in discrete (non-overlapping) generations. In each generation, each individual, independently, produces a random number of potential offspring according to a given law. From the pool of potential offspring N of them are then sampled uniformly at random and without replacement to survive to maturity and replace the parents. If there are fewer than N potential offspring from which to sample, we suppose that the population is unchanged over the generation (all the potential offspring perish before reaching maturity)*.

In Definition 2.3 we do not specify a law for the potential offspring. The key idea is that the reproducing individuals independently contribute potential offspring according to the same nontrivial law (i.e. the offspring numbers are not just independent but also identically distributed). Since we are only interested in models in which the total number of potential offspring produced at the same time exceeds the size (*N*) of the adult population with high probability (the mean number of potential offspring produced by any given individual exceeds 1) the specification of what happens when there are fewer than *N* potential offspring will be completely irrelevant. Schweinsberg’s model has been extended to diploid (where each individual carries two gene copies) populations, and the extension has been shown to give rise to Beta-coalescents admitting simultaneous multiple mergers (Birkner et al., 2018), in line with the results of Möhle and Sagitov (2003) and Sagitov (2003).

Before we can state the convergence theorem for the convergence to a Beta-coalescent from Schweinsberg’s model, we define a quantity called the coalescence probability.

#### Definition 2.4

(The coalescence probability *c*_*N*_). *Define c*_*N*_ *as the probability of the event that two individuals, chosen uniformly at random without replacement and at the same time from a given population derive from the same individual present in the previous generation*.

In Definition 2.4 the subscript *N* stands for population size, and it will be assumed that *c*_*N*_ *>* 0 for all *N* ∈ ℕ. In a haploid panmictic population of size *N* evolving according to the Wright-Fisher model, *c*_*N*_ = 1*/N*, and the ancestral process is the Kingman coalescent with time measured in units of *N* generations.

The concept of effective population size is a key concept in population genetics, and for our discussion, and is related to *c*_*N*_ (Definition 2.4). Throughout we take effective size as now stated (recall *c*_*N*_ *>* 0 for every *N* ∈ ℕ by assumption).

#### Definition 2.5

(Effective population size). *We define effective population size, denoted N*_*e*_, *as*

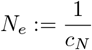

When the population evolves according to Definition 2.3 with law as in (1.1) for the number of potential offspring 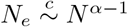 as *N* → ∞ when 1 *< α <* 2 (Schweinsberg, 2003).

#### Theorem 2.6

(Schweinsberg (2003)). *Suppose that a haploid population of size N evolves according to Definition 2.3. Let X be the random number of juveniles produced by an individual so that* (1.1) *holds for* α > *0 and a normalising constant C* > *0*. *Recall that* {*ξ* ^*n,N*^*(t),t ≥0* } *denotes the ancestral process for a sample of size n. Then*{ *ξ*^*n,N*^ (⌊ *t/c*_*N*_ ⌋), *t≥* 0 *converges, in the sense of convergence of finite-dimensional distributions, as N to a process whose law if α* 2 *is that of the Kingman coalescent restricted to* [*n*] *and if α* ∈ [1, 2) *is that of the Beta-coalescent (Definition 2.2) restricted to* [*n*].

Under the assumptions of Theorem 2.6, if 0 *< α <* 1 the ancestral process converges to a discrete time analogue of a Ξ-coalescent; with the measure Ξ determined by *α* and no scaling of time (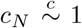; Schweinsberg, 2003). We do not consider this scenario here. The reason is that since mutation rates are ‘small’ (see Section 2.2), it is not easy to explain any observed genetic variation without a rescaling of time, meaning that we require sufficient time between merger events involving ancestors of the sample to see any mutations in the sample. For the same reason we also exclude the case *α* = 1 since then *c*_*N*_ log 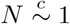(Schweinsberg, 2003).

### 2.2 Mutation and the site frequency spectrum

In practice, one does not observe the coalescent process (i.e. gene genealogies) directly, but rather a projection of it, determined by the mutations that fall on the coalescent tree (see Figure 1 for examples). We assume that the infinitely many sites (a ‘site’ refers to location on a chromosome) model of mutations (Kimura, 1969; Watterson, 1975) applies, so that an individual retains all the mutations that fall on its ancestral line (i.e. each new mutation occurs at a new site). For a sample of size *n*, we define the *unfolded site frequency spectrum* (abbreviated SFS) to be the random vector (*M*_1_(*n*), *M*_2_(*n*), …, *M*_*n*−1_(*n*)), where *M*_*j*_(*n*) is the random number of new (or derived, i.e. different from the ancestral state) mutations carried by exactly *j* of the *n* individuals (i.e. DNA sequences) in the sample. The expected value of *M*_*j*_(*n*) is the expected number of mutations to fall on branches of the genealogical tree that are ancestral to exactly *j* individuals in the sample. This shows that, although the SFS is a simple summary statistic of the full DNA sequence data, it contains valuable information about the pattern of observed genetic variation. Write 𝕞 for the mutation rate per locus per generation (by a *locus* we mean a contiguous non-recombining segment of a chromosome; a locus can consist of any number of sites) Under the Kingman coalescent, 𝔼 [*M*_*j*_(*n*)] = 2*N*_*e*_𝕞*/j* for *j* = 1, 2, …, *n*− 1 (Fu, 1995). For general Λ- and Ξ-coalescents there are numerical methods for computing 𝔼 [*M*_*j*_(*n*)] exactly (Birkner et al., 2013b; Blath et al., 2016; Spence et al., 2016).

**Figure 1:**
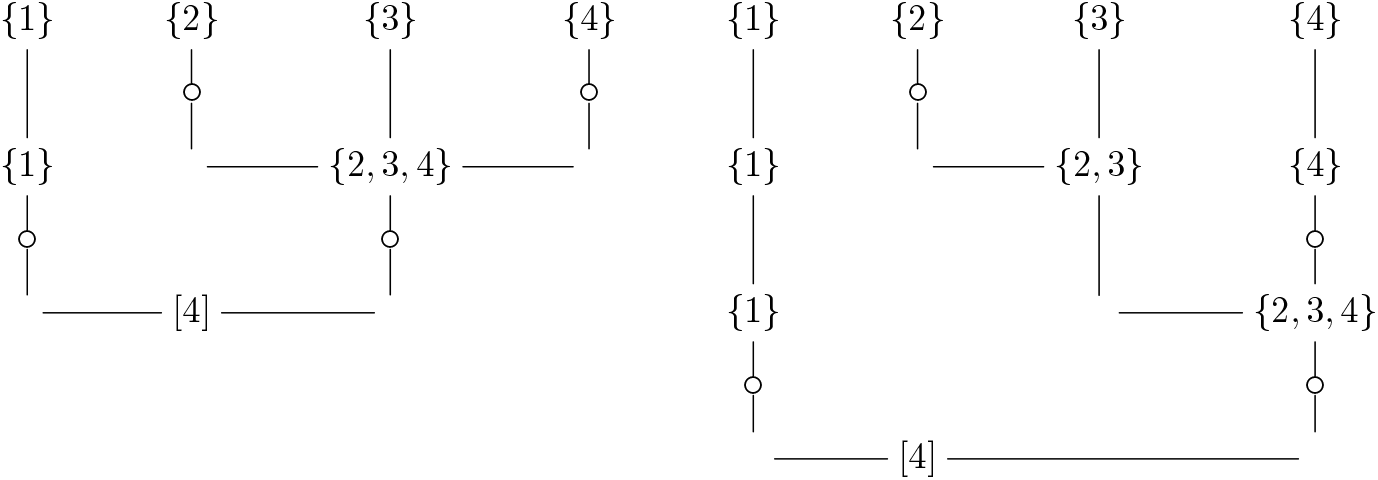
Two trees for *n* = 4, mutations are shown as ‘°’; the observed site-frequency spectra would not allow to distinguish between the two gene genealogies

The amount of information available to us about the genealogy of a sample is strictly limited by the mutation rate; it would have to be infinite for us to be able to resolve the ancestral tree of sampled gene copies entirely (see Figure 1 for an example). It is useful to quantify this. The quantity *N*_*e*_ is sometimes called the coalescent effective population size (Sjödin et al., 2004; Wakeley and Sargsyan, 2008). It is the amount by which time must be rescaled in order for a limiting coalescent to approximate the genealogy of a sample from a population evolving according to a given population model. In the (haploid) Schweinsberg model, if the limiting coalescent is the Beta-coalescent with parameter *α*∈ (1, 2), the corresponding scaling of time will be ⌊*AN* ^*α*−1^⌋ generations per coalescent time unit, where *A* = 𝔼 [*X*]^*α*^ */*(*αCB*(2 −*α, α*)), and *N* is the population size (Schweinsberg, 2003). In the Beta-coalescent, the expected time (measured in coalescent time units) until the first coalescence event among a sample of *n* lineages is of order 1*/n*^*α*^. Then if we are to see any mutations at all before the first merger, which is certainly not enough to resolve the coalescent tree, we require 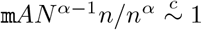 for large *N*; that is (at least) 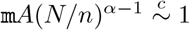. The mutation rate per site per generation has been estimated at around 2.2 × 10^−9^ in mammals (other than humans) (Kumar and Subramanian, 2002), in humans specifically around 1.2 × 10^−8^ (Kong et al., 2012), at 8.3 × 10^−9^ in Pacific cod (CANINO et al., 2010), and a study of the mtDNA of 32 Atlantic cod found 298 polymorphic sites over 15655 sites, and gave an estimated mutation rate of 1.1 × 10^−8^ (Carr and Marshall, 2008). From the mtDNA study on Atlantic cod, we see that the mutation rate per mtDNA genome per generation is approximately 10^−4^ (Carr and Marshall, 2008). For comparison with bacteria, Lee et al. (2012) estimated the spontaneous mutation rate in *E. coli* to be around 1 × 10^−10^ per nucleotide per generation, or around 1 × 10^−3^ per *E. coli* genome (around 4.6Mb) per generation. Given these mutation rate estimates, we see that the requirement 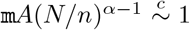 for large *N* becomes harder to fulfill as *α* moves closer to one, i.e. as the effect of sweepstakes reproduction increases. For example, suppose *X* ≥ 1, and ℙ [*X* ≥ *k*] = *C/k*^*α*^ for *k* ∈ {1, 2, …}, then *C* = 1, and 𝔼 [*X*] ≈ 1*/*(*α* − 1). Since *αCB*(2 − *α, α*) will be close to one when *α* approaches one, our requirement 𝕞*A*(*N/n*)^*α*−1^ ≥ 1, for a concrete example, is equivalent to

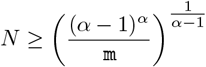

The exponent 1*/*(*α −*1) blows up the required population size to unrealistic values as *α →* 1. For an example, take *α* = 1.01 (Árnason and Halldórsdóttir, 2015), with 𝕞 ≈ 10^−5^ representing the approximate mutation rate over a 1 kb contiguous non-recombining chromosome segment in cod, to see that we would require *N* to be at least 10^300^ to recover the observed mutations used to estimate *α*.

### 2.3 Increasing sample size

Previous studies of the effect of increasing sample size (when sample size is at least proportional to *N*_*e*_ defined in Definition 2.5) on gene genealogies have focussed on the Wright-Fisher model (Wakeley and Takahashi, 2003; Fu, 2006; Bhaskar et al., 2014; Melfi and Viswanath, 2018a,b). When investigating the impact of such sample sizes on inference in population genetics, one might want to ‘calibrate’ the sample size against the effective size of the relevant population model. Wakeley and Takahashi (2003) consider a haploid population of constant size *N* evolving according to the Wright-Fisher model. In order to allow the sample size to exceed *N*_*e*_, they assume that in each generation *N −N*′ *>* 0 individuals die without reproducing, and then assign *N* offspring to *N*′ parents by multinomial sampling with equal weights. Clearly, (recall *c*_*N*_ from Definition 2.4) *c*_*N*_ = 1*/N*′, so *N*_*e*_ = *N*′. Through both analysis and simulation, they demonstrate that when sample size exceeds *N*_*e*_ one observes an excess of singletons (mutations observed in a single copy) in the SFS, compared to the predictions of the Kingman coalescent.

In Section 3.3, we give arguments for why in the Wakeley and Takahashi (2003) model the genealogy of a sample will, with high probability, deviate from the Kingman coalescent as soon as the sample size is of the order of 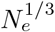. However, the deviation of the SFS from the predictions of the Kingman coalescent only becomes evident when the sample size is proportional to *N*_*e*_, at which point it reveals itself in part as an excess of singletons. We also recall (see e.g. Melfi and Viswanath (2018a) and Melfi and Viswanath (2018b)) an explanation for why we need such large sample sizes for the effect to become noticeable.

Wakeley and Takahashi (2003) re-analyse mitochondrial DNA data of introduced populations of Pacific oysters from Boom et al. (1994) and show that under their model, in order to explain the excess of singletons simply through large sample size relative to effective population size, one requires an implausibly large mutation rate. This might imply that sweepstakes reproduction is the true explanation for the excess. Multiple merger coalescents do not necessarily reflect a small effective population size compared to census population size, and a consequent loss of genetic diversity, and therefore may not require an implausibly high mutation rate to explain observed data (cf. e.g. Eldon (2020) and Remark 3.5).

## 3 Mathematical results

In this section we present the main mathematical results. We start by introducing an extension of (1.1) behind the main results.

### 3.1 The extended Schweinsberg model

We will be concerned with a population as described in Definition 2.3. We will consider an extension of the model considered by Schweinsberg (2003) (recall (1.1)); in our version the distribution of the number of potential offspring produced by each individual can depend on the total population size. We write *X*^*N*^ for the random number of potential offspring of an arbitrary individual in a population of size *N*, and we will write 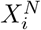 for the random number of potential offspring produced in a given generation by the *i*th individual in the population. The 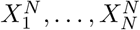 are independent and identically distributed (abbreviated i.i.d.) copies of *X*^*N*^, and the superscript *N* is to remind us that the range of *X*^*N*^ depends on *N* (which is not the case in the model considered by Schweinsberg (2003)). We will write

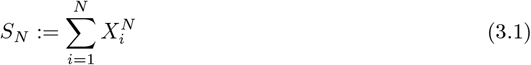

for the random total number of juveniles produced in any given generation. We suppose that 1 *< α <* 2 (see Remark 3.1), *ψ*(*N*) is a positive function of *N*, and there exist non-negative bounded functions *f* and *g* on ℕ, both independent of *N*, so that for all *k* ∈ [*ψ*(*N*)]

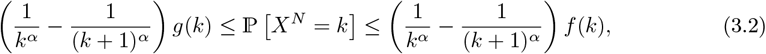

and we assign any remaining mass to {*X*^*N*^=0}. The quantity *ψ* (*N*) serves as an upper bound on the number of juveniles each individual can produce, i.e. ℙ [*X*^*N*^ ≤ *ψ*(*N*)] = 1. Since ℙ [*X*^*N*^ = *k*] = ℙ *[X* ≥ *k*] − ℙ [*X*^*N*^ ≥ *k* + 1] we see that ℙ[ *X*^*N*^ = *ψ*(*N*)] = ℙ [*X*^*N*^ ≥ *ψ*(*N*)] since ℙ [*X*^*N*^ ≥ *ψ*(*N*) + 1] = 0 by assumption. The model in (3.2) is an extension of the model in (1.1) in the sense of incorporating an upper bound on the number of potential offspring any given individual can produce. We will identify conditions on *f* and *g* so that the ancestral process will converge to a non-trivial limit; working with the mass function of the form ℙ [*X*^*N*^ = *k*] = (*k*^−*α*^ *−* (1 + *k*)^−*α*^)*h*(*k*) for a suitable *h* should lead to similar results.

We suppose that the following limits exist. With *g*_∞_, *f*_∞_, and *m*_∞_ denoting positive constants we take

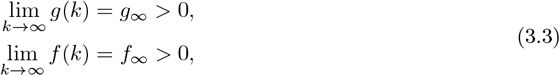

and, as *N* → ∞,

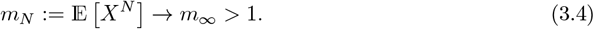

Recall from Definition 2.3 that if *S*_*N*_ *< N* then we simply maintain the original population over the generation. The functions *f* and *g* should be such that 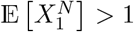 (and so 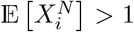 for all *i* ∈ [*N*] since the 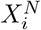 are taken to be i.i.d.). We define, for *k* ∈ ℕ,

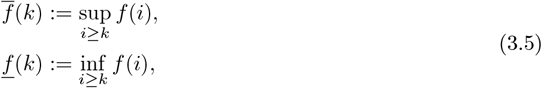

with 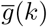and *g*(*k*) similarly defined. Then, for all *b* ∈ {1, …, *ψ*(*N*)},

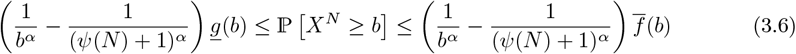

As *g*_∞_ *>* 0 by assumption (see (3.3)), *g* can be chosen to ensure *g*(*i*) *>* 0 for all *i* and we will assume this throughout. From (3.2) we see that, with 1 *< α <* 2,

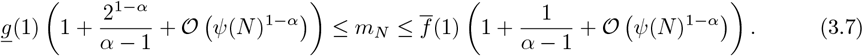

It is clear from (3.7) that if *g*(1) is big enough then *m*_*N*_ will be greater than 1. In Section 3.2 we discuss the units of time associated with the coalescents derived from our model.

### 3.2 Timescale and coalescents

For stating the timescaling associated with our model in (3.2) it will be convenient to define (cf. Lemma 6 in Schweinsberg (2003); recall *S*_*N*_ from (3.1))

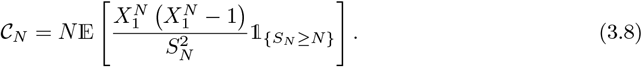

In Lemma 7.12 we prove that (*c*_*N*_ is defined in Definition 2.4)

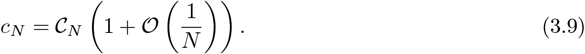

#### Remark 3.1.

*For α* ≥ 2 *we recover the Kingman coalescent from* (3.2) *regardless of ψ*(*N*). *The effect of large sample size on the approximation of the Kingman coalescent to gene genealogies of samples from a population evolving according to the Wright-Fisher model has been investigated (Wakeley and Takahashi, 2003; Melfi and Viswanath, 2018b,a; Bhaskar et al*., *2014; Fu, 2006). Our focus is on random sweepstakes and multiple-merger coalescents. Unless otherwise stated, Definition 2.3 is in force with numbers of potential offspring produced according to* (3.2) *restricted to* 1 *< α <* 2.

Our first result (Proposition 3.2) identifies 𝒞_*N*_ (see (3.8)) with error estimates and given that Definition 2.3 and (3.2) are in force. Most work in population genetics concerned with deriving coalescents ignores the error in the approximation. Controlling the order of the error in the coalescent approximations provides insight into the variability that one can expect in estimates of the effective population size under a given model. The effective population size is a key quantity in population genetics; estimating it has therefore received substantial consideration (Nei and Tajima, 1981; Felsenstein, 1992; Hill, 1981; Waples, 2006; Beerli and Felsenstein, 1999; Fu, 1994; Berthier et al., 2002; Palstra and Ruzzante, 2008; Palstra and Fraser, 2012; Hedgecock, 1994a). Identifying the sources of error in the coalescent approximation might help with understanding deviations from expectations, and designing sampling and DNA sequencing strategies (Felsenstein, 2005; Pluzhnikov and Donnelly, 1996). Small effective relative to census population size has been interpreted as evidence of sweepstakes reproduction (Waples, 2016). We will show that there can be large differences in predictions of the effective size, even when converging to the same coalescent, and that, in particular in our model, the asymptotic behaviour of *ψ*(*N*)*/N* plays a key role. Section 7 contains a proof of Proposition 3.2.

#### Proposition 3.2.

*Suppose a haploid population evolves according to Definition 2.3 and* (3.2) *with* 1 *< α <* 2. *Let K* > 0 *be a constant, recalling* (3.3) *suppose f*_∞_ = *g*_∞_, *recall* 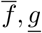, *g from* (3.5), *m*_*N*_ *from* (3.4), *and* 𝒞_*N*_ *from* (3.8). *In each of the statements below, L is a function of N, over which one can optimise for any specific choice of model (see Section 7, e*.*g. Lemma 7.4); the quantities* 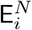 *denote the errors in our estimates*.

1. *Suppose that ψ*(*N*)*/N* → 0, *and L*(*N*)*/ψ*(*N*) → 0. *Then*

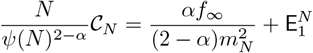

*where (recall* (7.6) *and Remark 7.9 with η* = (*α* + 1)*/*(2*α*))

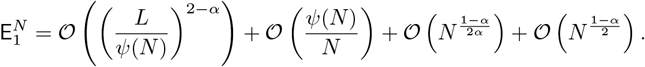
2. *Suppose that ψ*(*N*)*/N* → *K, and L*(*N*)*/N* → 0. *Then*

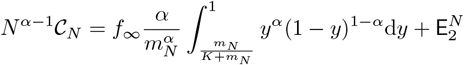

*where (recall* (7.6) *and Remark 7.9 with η* = (4 − *α*)*/*3)

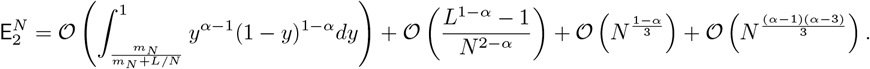
3. *Suppose that ψ*(*N*)*/N* → ∞ *and L*(*N*)*/N* → 0. *Then*

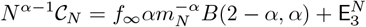

*where (recall* (7.6) *and Remark 7.9 with η* = 2*/*(1 + *α*))

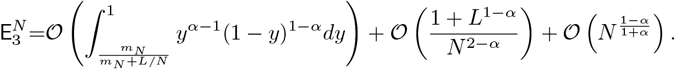

The constant term in Case 3 in Proposition 3.2 (that is the limit as *N* → ∞ of *N* ^*α*−1^ 𝒞_*N*_) is the same as that obtained in Lemma 13 of Schweinsberg (2003) where there is no restriction on the number of potential offspring. Further, 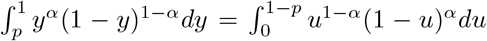 for any 0 ≤ *p* ≤ 1.

Before discussing the implications of Proposition 3.2, we state two more propositions, which detail the limiting coalescents for the three cases in Proposition 3.2. We identify the limiting coalescent for fixed sample sizes, again maintaining control over the errors. Section 9 contains a proof of Proposition 3.3.

#### Proposition 3.3.

*Under the conditions of Proposition 3.2 Case (1), the scaled ancestral process* {*ξ*^*n,N*^ (⌊*t/c*_*N*_ ⌋), *t* ≥ 0} *corresponding to a sample of size n converges to the Kingman coalescent restricted to* {1, …, *n*}. *Moreover (recall* (2.3) *in Definition 2.1) in a large population*

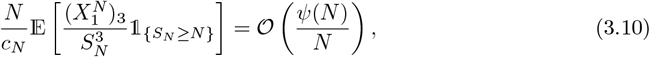

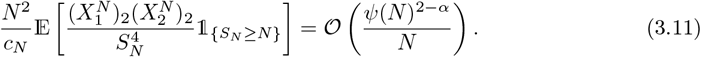

We now specify the limiting coalescent for the cases in which multiple-merger coalescents are obtained. Section 10 contains a proof of Proposition 3.4.

#### Proposition 3.4.

*Under the conditions of Proposition 3.2 Case 2, the scaled ancestral process* {*ξ*^*n,N*^ (⌊*t/c*_*N*_ ⌋), *t* ≥ 0} *corresponding to a sample of size n converges to a process whose law is given by a* Λ*-coalescent (see* (2.6)*) restricted to* {1, …, *n*} *and without an atom at zero (corresponding to a* = 0 *in* (2.6)*), with m*_∞_ *defined in* (3.4) *and K* > 0 *fixed. The* Λ_+_ *measure in* (2.6) *is given by*

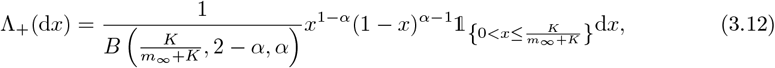

*where we have used the notation (recall B*(*p, a, b*) *from* (2.5))

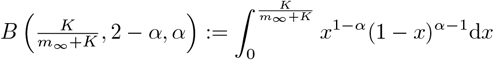

*Under the conditions of Proposition 3.2 Case 3, the scaled ancestral process* {*ξ*^*n,N*^ (*t/c*_*N*_), *t* ≥ 0} *corresponding to a sample of size n converges to a process whose law is given by a* Λ*-coalescent restricted to* {1, …, *n*}, *with the* Λ*-measure given in* (2.7). *Moreover, in Cases 2 and 3, as N* → ∞,

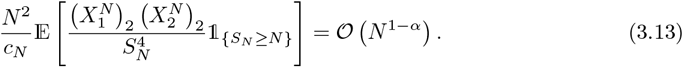

#### Remark 3.5.

*In Section 2.2 we discuss that for α close to one it may be difficult to recover the observed amount of genetic variation for a given sample with the timescale of the multiple-merger coalescents we are considering. To overcome this, we can take ψ*(*N*) *to be random. Suppose* (*ψ*_1,*N*_)_*N*∈ℕ_ *and* (*ψ*_2,*N*_)_*N*∈ℕ_ *are sequences of positive numbers, where ψ*_1,*N*_ */N* → 0, *and ψ*_2,*N*_ */N* ≩ 0, *where ψ*_2,*N*_ */N* ≩ 0 *means ψ*_2,*N*_ */N* > 0 *for all N, and* lim inf_*N*→∞_ *ψ*_2,*N*_ */N* > 0. *Take*

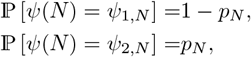

*independently in each generation. To clarify, in every generation the* 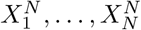 *are i*.*i*.*d*., *so ψ*(*N*) *takes the same value for all* 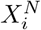 *in each generation, but ψ*(*N*) *may vary between generations as just described. If p*_*N*_ = (*ψ*_1,*N*_ */N*)^2−*α*^ *then* 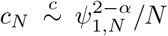 *and the scaled ancestral process converges to a* Λ*-coalescent which is a mixture between Kingman and Beta-coalescent (cf*. (2.6)*). This is an easy consequence of the calculations in Sections 9 and 10 and Proposition 6.2. The coalescent rate for the merging of k out of n blocks* (2 ≤ *k* ≤ *n*) *is then of the general form, with* 0 *< γ* ≤ 1 *and c, c*′ > 0 *all fixed*

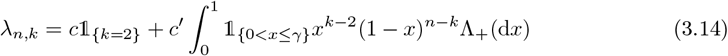

*where* Λ_+_ *has no atom at zero (recall Proposition 3.4). The new timescaling (proportional to N generations whenever ψ*_1,*N*_ = *O*(1), *taking ψ*_1,*N*_ = *N/* log *N leads to* 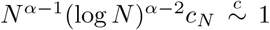*) means that lower mutation rates are required to explain the observed genetic diversity*.

Propositions 3.2-3.4 give us an insight into the sources of the errors in the coalescent approximation. An uncertainty in the estimate of the effective size can come through the scaling constant by either assuming a wrong model for the upper bound (*ψ*(*N*)), or if *ψ*(*N*) varies considerably over the ancestral history of our sample. A further source of error can come from ignoring the error terms.

Case 1 in Proposition 3.2 reveals that the timescale and some of the error terms depend on *ψ*(*N*), even though the limiting process is the Kingman coalescent. Suppose *ψ*(*N*) = *N/* log *N*. Then *ψ*(*N*)*/N* → 0 so we would be in the domain of attraction of the Kingman coalescent, but time would be measured in units proportional to *N* ^*α*−1^(log *N*)^2−*α*^ generations, so we would be measuring time in essentially the same way as for the Beta-coalescents. Thus, even when we are in the domain of attraction of the Kingman coalescent, the effective size can be small relative to *N*. The effective size can range from *cN* to *c*′*N* ^*α*−1^(log *N*)^2−*α*^ for some *c, c*′ > 0 fixed and with 1 *< α <* 2. Considering the error terms, two of them are of order *N* ^1−*α*^, and this can be arbitrarily close to 𝒪 (1). Further, with *ψ*(*N*) = *N/* log *N* we see that the error term of order *ψ*(*N*)*/N* decreases only as 1*/* log *N*.

For both Case 2 and Case 3, the leading term decreases as *α* → 1. The error terms of the form *N* ^*c*(1−*α*)^ for some *c* > 0 can be arbitrarily close to 𝒪 (1). The uncertainty in estimating effective size can come from two directions; in Case 1 from uncertainty in determining the order of 𝒞_*N*_, and in all three cases from the (at least potentially) large errors. Since 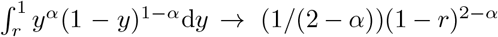 as *r* → 1, we see 1*/*𝒞_*N*_ for Case 2 can be much larger than 1*/*𝒞_*N*_ for Case 3 if *K* is much smaller than *m*_*N*_.

Propositions 3.3 and 3.4 tell us that when 1 *< α <* 2 (see Remark 3.1) it is the behaviour of *ψ*(*N*)*/N* (as *N* → ∞;) that determines the limiting process. We obtain a Kingman coalescent in case *ψ*(*N*)*/N* → 0, and a multiple merger coalescent when *ψ*(*N*)*/N* ≩ 0, i.e. the limiting measure Λ is of the form

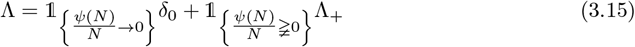

where Λ_+_ is a finite measure on (0, 1] (without an atom at zero), and *ψ*(*N*)*/N* ≩ 0 is as defined in Remark 3.5.

### 3.3 Increasing sample size and the Kingman coalescent

We wish to establish how large a sample size we can take before the errors that we make in using the limiting coalescent that we have identified to estimate the distribution of the site frequency spectrum become significant. Since mutation rates are not infinite, statistics based on the SFS will not, in particular, be able to differentiate between mergers that happen instantaneously and those which occur in quick succession, if both lead to the same topology for the genealogical tree.

First, we consider the impact of large sample size on genealogies for two population models in the domain of attraction of the Kingman coalescent (Sections 3.3.1 and 3.3.2).

#### 3.3.1 The Wakeley-Takahashi model

We begin with a brief discussion of a variant of the Wright-Fisher model, that we will refer to as the Wakeley-Takahashi model (Wakeley and Takahashi, 2003). We will give arguments for the result, that if the sample size *n* is at least proportional to *N*_*e*_^1*/*3^, where *N*_*e*_ is the effective size (recall Definition 2.5) in the Wakeley-Takahashi model, one can expect to see multiple-mergers in the genealogy. We will also argue, that one can only expect to see effects of increasing *n* on the SFS when *n* is at least proportional to *N*_*e*_.

In the Wakeley-Takahashi model, a haploid panmictic population of constant size *N* evolves in discrete generations. In each generation, first sample *N*′ ≤ *N* ‘potential’ parents, uniformly at random, from the population; and then determine the family sizes of those *N*′ individuals by multinomial sampling with equal weights. In other words, each of the *N* offspring chooses a parent uniformly at random with replacement from among the subset of *N*′ available to reproduce. Thus, after the first generation, the Wakeley-Takahashi model reduces to the Wright-Fisher model with population size *N*′. It then holds that *N*_*e*_ = *N*′. This construction allows one to ask what happens to the genealogy when *N*_*e*_ ≤ *n* ≤ *N*, i.e. when the sample size is at least equal to the effective size.

When *N*_*e*_ = *N* a detailed analysis of the coalescent for large samples is provided by Melfi and Viswanath (2018a). Now we give arguments for the 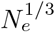 result. Two types of events can cause deviations from the predictions of the Kingman coalescent: *(i)* asynchronous and *(ii)* simultaneous multiple mergers (where a multiple merger involves at least three lineages). The probability of seeing two distinct pairs of lineages merge into separate parents in a single generation when there are *k* lineages is (recalling (2.3)) 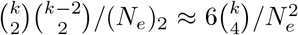. Similarly, the probability of seeing a merger of three lines in a single generation when there are *k* lineages is 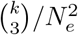. If neither of these events happens, the number of generations that it takes for the number (*k*) of ancestral lineages to change is about 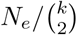 (on average), so the chance that we see a simultaneous merger before we see our first pairwise (involving two lineages) coalescence event is (up to a combinatorial constant that does not depend on *k*) approximately 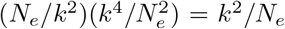. To estimate the chance that we see a simultaneous merger before the most recent common ancestor of a sample of size *n*, we sum the terms *k*^2^*/N*_*e*_ over *k* = 2, …, *n* (recall that 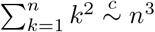 as *n* → ∞) to find that the probability is proportional to *n*^3^*/N*. In other words, if *n* is at least proportional to 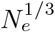 there is an appreciable probability that we will see a simultaneous merger of two pairs of lineages somewhere in the genealogical tree.

Although these arguments suggest that we can expect the genealogy of a sample to differ from that predicted by the Kingman coalescent once *n* is at least proportional to 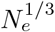, the results of Wakeley and Takahashi (2003) illustrate that this is not evident in the site-frequency spectrum. Indeed, one only starts to see significant deviation of the SFS from the predictions of the Kingman coalescent when *n* is proportional to *N*_*e*_ (Wakeley and Takahashi, 2003). It has been observed by Bhaskar et al. (2014) that simultaneous pairwise mergers may not be too disruptive. To see why, notice that when the number of ancestral lineages is large, and so coalescence events are frequent, we can only hope to detect the difference between two simultaneous mergers (event A in Figure 2), and two mergers in quick succession, if those mergers lead to different tree topologies; that is if, in the second scenario, the parental lineage of the first merger is also involved in the subsequent merger (event B in Figure 2).

**Figure 2:**
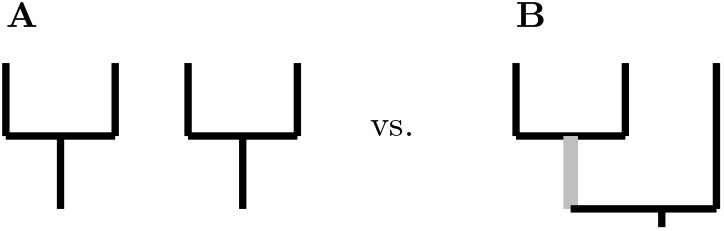
Illustration of events leading to different tree topologies

If there are currently *k* ancestral lineages, event B happens with probability approximately 1*/k*, and our heuristic above says that we will see such events with a frequency of the same order as triple mergers. Even if this happens, the SFS is only likely to be changed if such anomalous events make up a significant proportion of coalescence events; that is they must occur on a comparable timescale to simple pairwise mergers. This requires 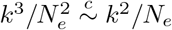 as *N*_*e*_ increases, which in turn requires sample sizes proportional to *N*_*e*_. This agrees with the results of Melfi and Viswanath (2018b), who compare the Wright-Fisher site-frequency spectrum with that predicted by the Kingman-coalescent, and show that one only starts to notice discrepancies between the two when *n* is at least proportional to *N*_*e*_.

We close this section with an argument for why we expect to see an increase in singletons with increasing sample size under the Wakeley-Takahashi model with effective size (recall Definition 2.5) *N*_*e*_. Once the size of a sample from a population evolving according to the Wakeley-Takahashi model with effective size *N*_*e*_ is significantly larger than *N*_*e*_, a lot of the external branch length arises from reducing *n* to something that is at most *N*_*e*_ over a single generation, and indeed if *n* increases too quickly relative to *N*_*e*_ then these will be the only external branches. To investigate this a bit further, if we distribute *n* individuals among *N*_*e*_ parents, then the expected number of parents with no offspring is *N*_*e*_(1 − 1*/N*_*e*_)^*n*^ *≈ N*_*e*_ exp(−*n/N*_*e*_) = 1 whenever *n* = *N*_*e*_ log *N*_*e*_ (understood to be ⌊*N*_*e*_ log *N*_*e*_⌋). If *n* = *bN*_*e*_ log *N*_*e*_ for any *b* > 1, then by Markov’s inequality, for large *N*_*e*_ with high probability, every one of the *N*_*e*_ parents has at least one offspring (with *ν* denoting the random number of offspring of an arbitrary individual we have ℙ [*ν ≥* 1] = 1 − ℙ [*ν* = 0] ≈ 1 − exp(− *n/N*_*e*_) = 1 − 1*/N*_*e*_ when *n* = ⌊*N*_*e*_ log *N*_*e*_ ⌋). Now think of distributing at least 2*bN*_*e*_ log *N*_*e*_ lineages - there is at least one of the first *bN*_*e*_ log *N*_*e*_ assigned to each of the *N*_*e*_ parents, and then at least one of the second *bN*_*e*_ log *N*_*e*_ assigned to each parent. Thus (with high probability) every family has at least two offspring and so it is only the initial generation during which *n* is reduced to at most *N*_*e*_ that contributes to the external lineages. Further discussion on this may be found in Appendix A. Another approach to identifying the sample size required to disrupt convergence to the Kingman coalescent is contained in Appendix B.

#### 3.3.2 The extended Schweinsberg model for the Kingman case

In this section we consider large sample size in connection with the extended Schweinsberg model as given in Definition 2.3 and (3.2) assuming *ψ*(*N*)*/N* → 0 (recall Proposition 3.3). We give arguments for the result, that a sample size proportional to 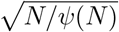 can be sufficient for the Kingman coalescent to be a poor approximation of the genealogy. For the extended Schweinsberg model 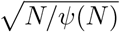 can be much smaller than 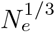 (recall Definition 2.5), the reference point for the Wakeley-Takahashi model (see Section 3.3.1). For an example, compare 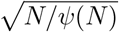 and (*N/ψ*(*N*)^2−*α*^)^1*/*3^ when *ψ*(*N*) = *N/* log *N*.

Even when the limiting coalescent is a Kingman coalescent (Case 1 in Proposition 3.2 where *ψ*(*N*)*/N* → 0), the extended Schweinsberg model that we consider here (see (3.2)) behaves rather differently from the Wakeley-Takahashi model. Recall *S*_*N*_ from (3.1), and (2.2) in Definition 2.1. Under the conditions of Proposition 3.3, when there are *k* lineages, we see simultaneous mergers at rate (relative to the rate of pairwise mergers; recall (3.11)),

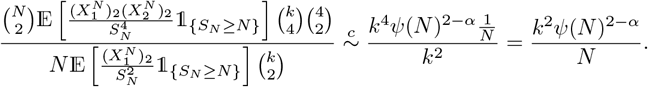

This is of order *k*^2^*/N*_*e*_ (cf. Case 1 in Proposition 3.2), just as in the Wakeley-Takahashi model. Summing over *k* = 2, …, *n* gives, as before, that we can expect to see simultaneous mergers in the genealogy of a sample of size *n* when *n* is at least proportional to 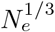.

On the other hand, we see 3-mergers at rate (relative to the rate of pairwise mergers; recall (3.10))

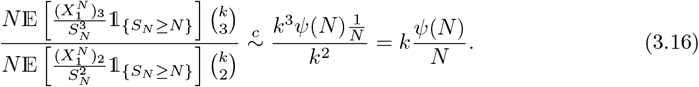

Summing as before gives, that we can expect to see 3-mergers in the genealogy when the sample size (where *c* > 0 is fixed)

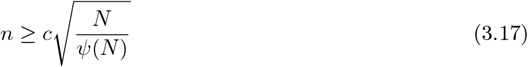

Suppose *N* ^1*/*(2*α*−1)^*/ψ*(*N*) → 0 and that *ψ*(*N*)*/N* → 0 (and recall 1 *< α <* 2). Then 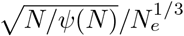 (recall Case 1 of Proposition 3.2) so we expect to see 3-mergers at a smaller sample size than simultanous pairwise mergers, so the number of lineages is going down one at a time and the summation over *k* from 2 to *n* is justified. When *N* ^1*/*(2*α*−1)^*/ψ*(*N*) ≫ 0 then 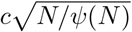 is an upper bound for the sample size at which we expect to see 3-mergers.

If *ψ*(*N*) is unbounded (but with *ψ*(*N*)*/N* → 0), this is significantly bigger than the 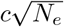 that we saw in the Wakeley-Takahashi model. This might not be surprising, but it suggests that the Kingman approximation for this model will be poor at sample sizes closer to 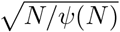, which could be substantially smaller than (*N/ψ*(*N*)^2−*α*^)^1*/*3^ (see Proposition 3.2, Case 1). As an example, take *ψ*(*N*) = *N/* log *N* to see that a sample size proportional to 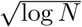 would then be sufficient for a poor approximation of the Kingman coalescent.

Suppose *ψ*(*N*) = *N*. Then we are in the domain of attraction of the incomplete Beta-coalescent, so that 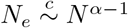 (recall Case 2 of Proposition 3.2); arguing as before then *N* ^(*α*−1)*/*3^ is an upper bound for the sample size at which we expect to see simultaneous mergers.

### 3.4 Increasing sample size and Beta-coalescents

In this section we focus on the Schweinsberg model (Definition 2.3) in the domain of attraction of the Beta-coalescents (recall (2.7) and (3.12)). In particular, we show that as a result of approximating sampling without replacement by sampling with replacement we consistently overestimate the rate of *k*-mergers (a merger of *k* lineages) and that the error is on the same order as the rate of mergers when *k* is proportional to 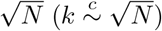). In Section 8.1 we provide a heuristic argument, based on known results for the number of blocks in the limiting coalescent, that suggests that this should significantly distort our prediction of statistics based on the site frequency spectrum for sample sizes of order *N* ^*α/*2^.

First we present a lemma which is similar in nature to Proposition 7.6, and from which the result 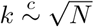 follows. From (3.18) in Lemma 3.6 one sees that, one would choose *L* (see Section 8 for the role of *L*) so that the maximum of (*L/N*)^*k*−*α*^ and *k*^2^*/L* is as small as possible. In a similar way one would optimise over *β*_*N*_. Section 8 contains a proof of Lemma 3.6.

#### Lemma 3.6.

*Consider a sample of n individuals from a population evolving according to Definition 2.3 and* (3.2) *with* 1 *< α <* 2. *Let* (*β*_*N*_)_*N*∈ℕ_ *be an increasing positive sequence. Suppose that n/N* → 0, *n* ≥ *k* ≥ 2, *f*_∞_ = *g*_∞_ *with f*_∞_ *and g*_∞_ *from* (3.3), *m*_*N*_ *from* (3.4), *S*_*N*_ *from* (3.1), *K* > 0 *a constant, and suppose L is a function of N with L/N* → 0.

1. *If ψ*(*N*)*/N* → *K*,

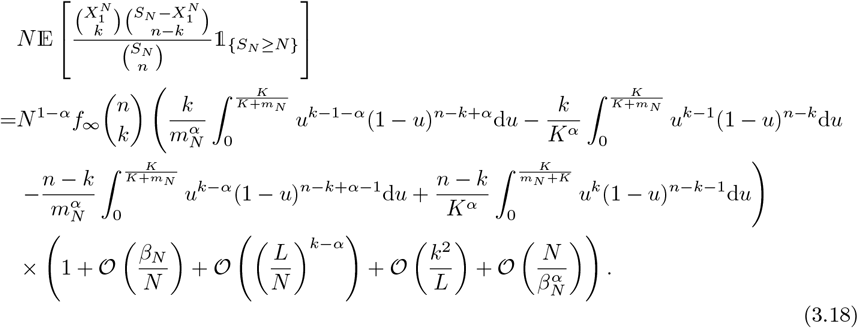
2. *If ψ*(*N*)*/N* → ∞,

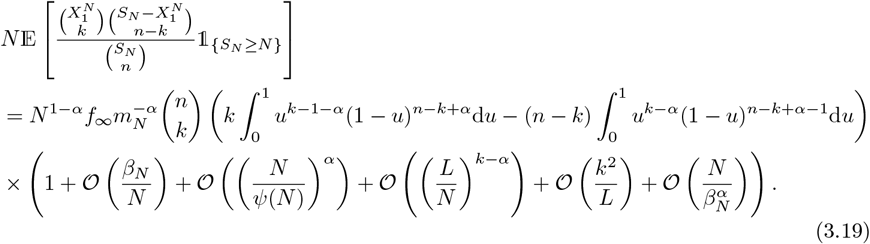

Equation (3.19) arises naturally in our proof and so we have left it in this form. However, observe that (recalling Γ(*x* + 1) = *x*Γ(*x*))

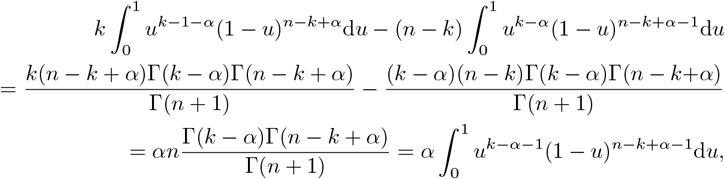

as expected from Proposition 3.4.

#### Corollary 3.7.

*When there are at least* 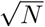 *lineages, then for large N we will not see convergence to the limiting rates suggested by Proposition 3.4*.

In particular, we see from the proof of Lemma 3.6 (see Section 8) that the 𝒪 (*k*^2^*/L*) error term is always negative. Therefore, when the limiting Λ-coalescent has mergers of at least 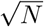 lineages we will be seeing a reduced number of such mergers, and so a different topology in the coalescent tree corresponding to the gene genealogy of the ancestral process. In Section 8.1 we present heuristic calculations to establish that a sample size proportional to *N*^*α/*2^ is sufficient for there to be an 𝒪 (1) probability of seeing a merger of at least 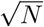 lineages in the limiting complete Beta(2 − *α, α*) coalescent tree (Definition 2.2). However, we will see in the next section that the site-frequency spectrum is not distorted by increasing sample size when the limiting tree is that predicted by the Beta coalescents considered here.

## 4 Increasing sample size and relative branch lengths

In this section we use simulations to investigate if and how increasing sample size may distort the site-frequency spectrum relative to the predictions of a given coalescent.

For the numerical examples we apply a special case of the model in (3.2). Consider a haploid population of fixed size *N* and evolving according to Definition 2.3. Let 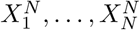 denote the independent and identically distributed random number of potential offspring produced in a given generation, where (recall [*n*] = {1, 2, …, *n*} for all *n* ∈ ℕ)

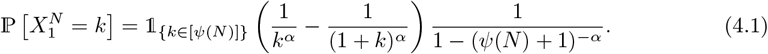

Then the mass function in 4.1 is monotone decreasing on [*ψ*(*N*)], i.e. 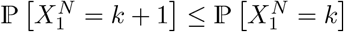. Suppose *K* > 0 is a constant, *ψ*(*N*)*/N* ≩ 0 (recall Remark 3.5), and write

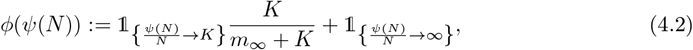

We will approximate *m*_∞_ (see (3.4) and (3.7)) with

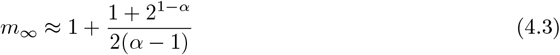

Our calculations then show that one obtains, with 1 *< α <* 2 and *ϕ* as in (4.2), a Beta coalescent with Λ_+_-measure given by (3.12) with *K/*(*m*_∞_ + *K*) replaced with *ϕ*. We also consider an unbounded distribution of number of juveniles (see Figure 4b), where *X* denotes the number of juveniles produced by an arbitrary individual, and

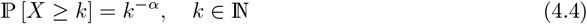

To describe the simulation results we need to define notation. Let *B*_*i*_(*n*) denote the random total length of branches supporting *i* ∈ [*n* − 1] leaves when the sample size is *n* and the gene genealogy of the sample is described by a given coalescent {*ξ*^*n*^}; let 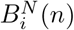 denote the corresponding quantity for a given ancestral process {*ξ*^*n,N*^}. With #*A* denoting the number of elements in a given (finite) set *A*, for all *i* ∈ [*n* − 1],

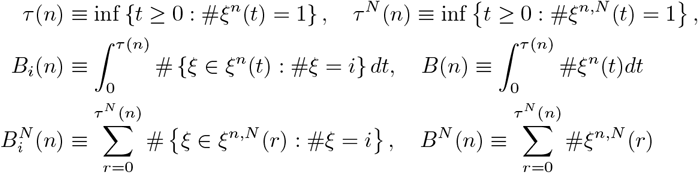

The branch lengths 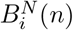 are recorded in generations. We are interested in understanding how increasing sample size may distort the site-frequency spectrum. To do so we sample gene genealogies from a finite population for increasing sample sizes, and compare estimates of mean relative branch lengths to the ones predicted by a given coalescent. Write, for *i* ∈ [*n* − 1],

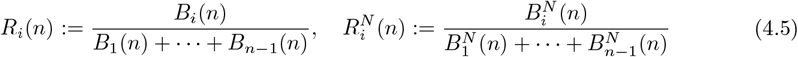

The quantities *R*_*i*_(*n*) and 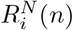 are well defined, since 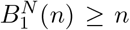, and *B*_1_(*n*) > 0 a.s. Write 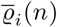 resp. 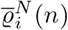 for the estimates of 𝔼 [*R*_*i*_(*n*)] resp. 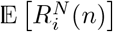.

We will compare 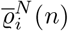 and 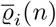. The 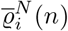 are obtained for samples from a finite haploid population evolving according to Definition 2.3 with numbers of potential offspring distributed according to (4.1) or (4.4).

In Figure 3 we compare 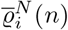 (symbols) and 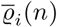 (black lines) when {*ξ* ^*n,N*^*}* is in the domain of attraction of the Kingman-coalescent. In Figure 3a we have taken *ψ*(*N*) = *N* and *α* = 3 so the effective size (recall Definition 2.5) is proportional to *N*. In Figure 3b we have taken *ψ*(*N*) = *N/* log *N* and *α* = 1.05, so the effective size is proportional to *N* ^*α*−1^(log *N*)^2−*α*^ (recall Case 1 of Proposition 3.2), and *N* ^*α*−1^(log *N*)^2−*α*^ *≈* 13 for *N* = 10^4^ and *α* = 1.05. The agreement between 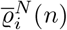 and 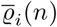 depends on the relation between the sample size and the effective size when the ancestral process is in the domain of attraction of the Kingman-coalescent. This is in line with results based on the Wright-Fisher model (Wakeley and Takahashi, 2003; Melfi and Viswanath, 2018a).

**Figure 3:**
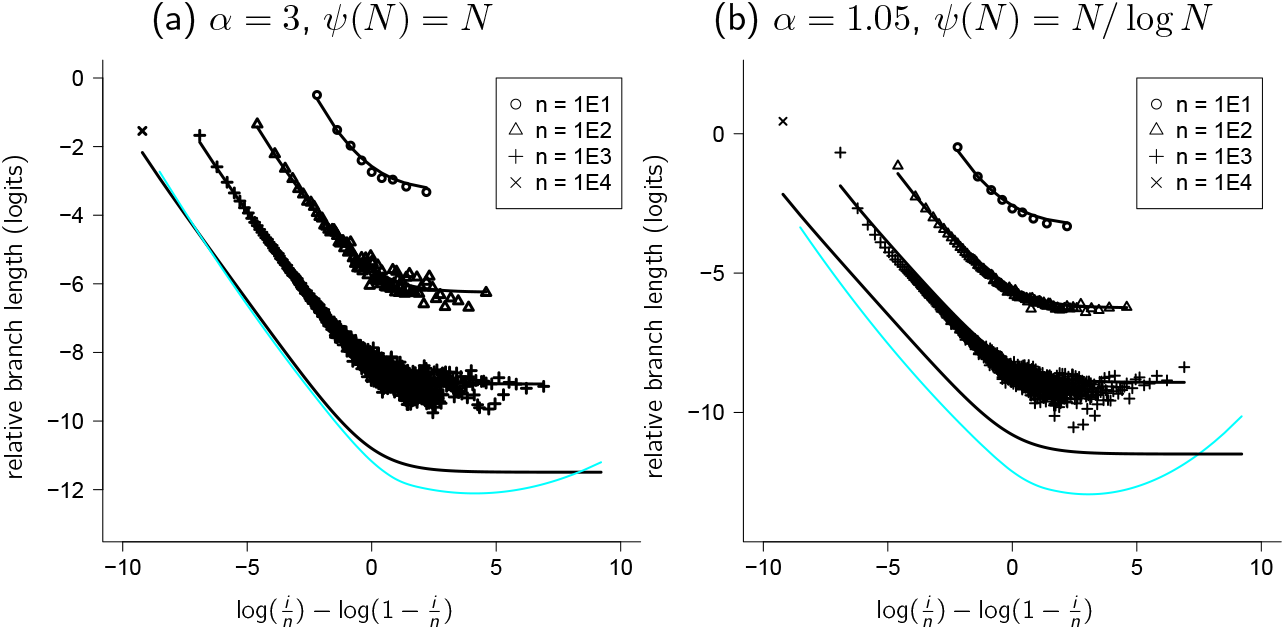
Comparing relative branch lengths – the Kingman coalescent. Comparison of 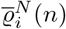 (symbols) and of 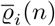 (black lines) shown as log(*e*_*i*_(*n*)) − log(1 − *e*_*i*_(*n*)) as a function of log(*i/n*) − log(1 − *i/n*) where *e*_*i*_(*n*) denotes the corresponding mean relative branch length estimate for sample size *n* as shown from a haploid panmictic population of constant size *N* = 10^4^ evolving according to Definition 2.3 with number of potential offspring distributed according to (4.1) with *α* and *ψ*(*N*) as shown. The black lines are the approximation 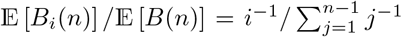 of 𝔼 [*R*_*i*_(*n*)] predicted by the Kingman coalescent (Fu, 1995). The cyan lines for the case *n* = *N* is a loess regression (using the function `loess in R` (R Core Team, 2023)) through 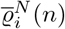 for *i* ∈ {2, 3, …, *n*−1}; the estimates of 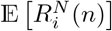 are the results from 10^4^ experiments. Appendix E contains a brief description of an algorithm for estimating 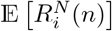

**Figure 4:**
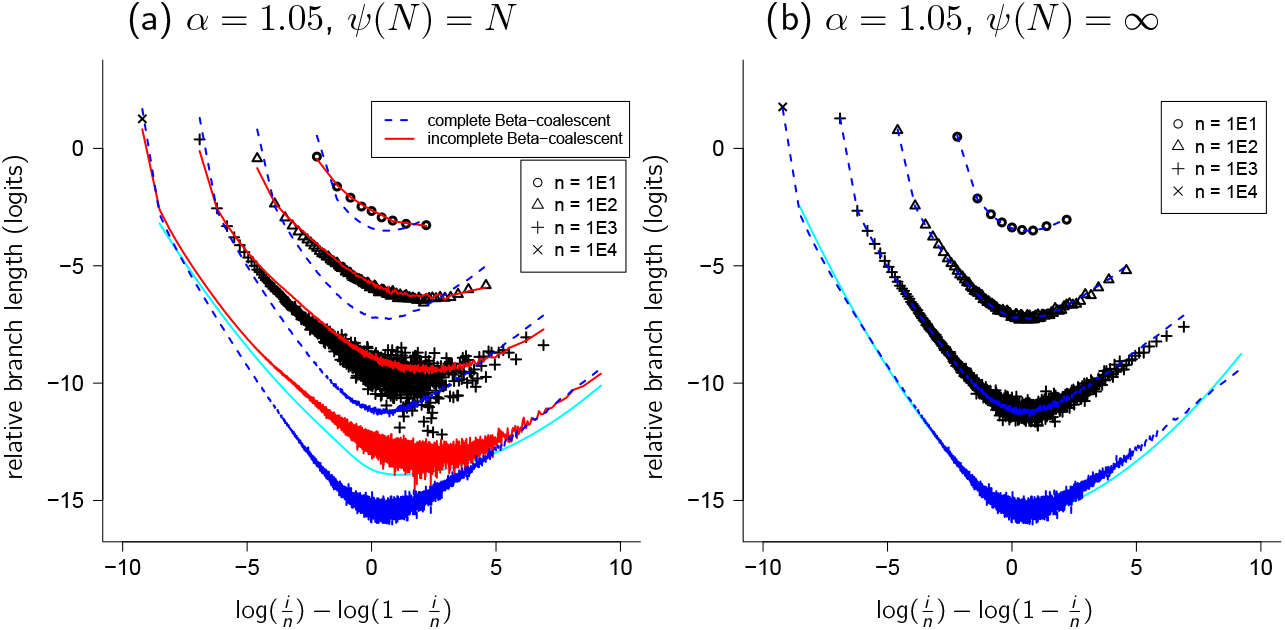
Comparing relative branch lengths - Beta-coalescents. Comparison of 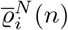 (symbols) and of 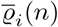 (lines) shown as log(*e*_*i*_(*n*)) − log(1 − *e*_*i*_(*n*)) as a function of log(*i/n*) − log(1 − *i/n*), where *e*_*i*_(*n*) denotes the corresponding mean relative branch length estimate for sample size *n* as shown from a haploid population of constant size *N* = 10^4^ evolving according to Definition 2.3 with number of potential offspring distributed according to (4.1) with *α* = 1.05 and *ψ*(*N*) = *N* (a), and according to (4.4) with *α* = 1.05 (b). The blue dashed lines show 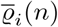 for the complete Beta-coalescent, and the red solid lines in (a) show 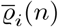 for the incomplete Beta-coalescent with *K* = 1. The cyan lines for the case *n* = *N* is a loess regression (using the function loess in R (R Core Team, 2023)) through 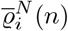 for *i* ∈ {2, 3, …, *n* − 1}; the estimates of 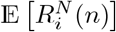 are the results from 10^4^ experiments. Appendix E contains a brief description of an algorithm for estimating 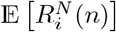

In Figure 4 we compare 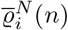 (symbols) and 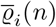 (blue and red lines) when *ξ* is in {*ξ* ^*n,N*^} is in the domain of attraction of the incomplete Beta-coalescent ((4.1); Figure 4a) and the complete Beta-coalescent ((4.4); Figure 4b). In Figure 4 the blue dashed lines are for the complete Betacoalescent, and the red solid lines in Figure 4a for the incomplete Beta-coalescent with *K* = 1. Overall the agreement between 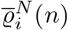 and 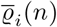 as predicted by the corresponding Beta-coalescent is good, indicating that the site-frequency spectrum predicted by our model is not distorted by increasing sample size when in the domain of attraction of a Beta coalescent. Figure 4a further shows that the estimates of 𝔼 [*R*_*i*_(*n*)] for the complete Beta-coalescent (blue dashed lines) do not match the estimates of 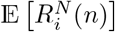 when *ψ*(*N*) = *N*, in contrast to the incomplete Beta-coalescent (red solid lines).

In Appendix C we compare relative branch lengths predicted by the Beta-coalescents (see Figure 5). The red line in Figure 5 is for the complete Beta(2−*α, α*) coalescent, and the remaining ones (except the black one is for the Kingman coalescent) are for the incomplete Beta(*γ*, 2 − *α, α*) coalescent with *γ* = *K/*(*K* + *m*_∞_) with *K* as shown. Figure 5 shows that incorporating an upper bound on the number of potential offspring can markedly affect the predicted site-frequency spectrum; the Beta-coalescent lines are all for the same *α* (*α* = 1.01).

**Figure 5:**
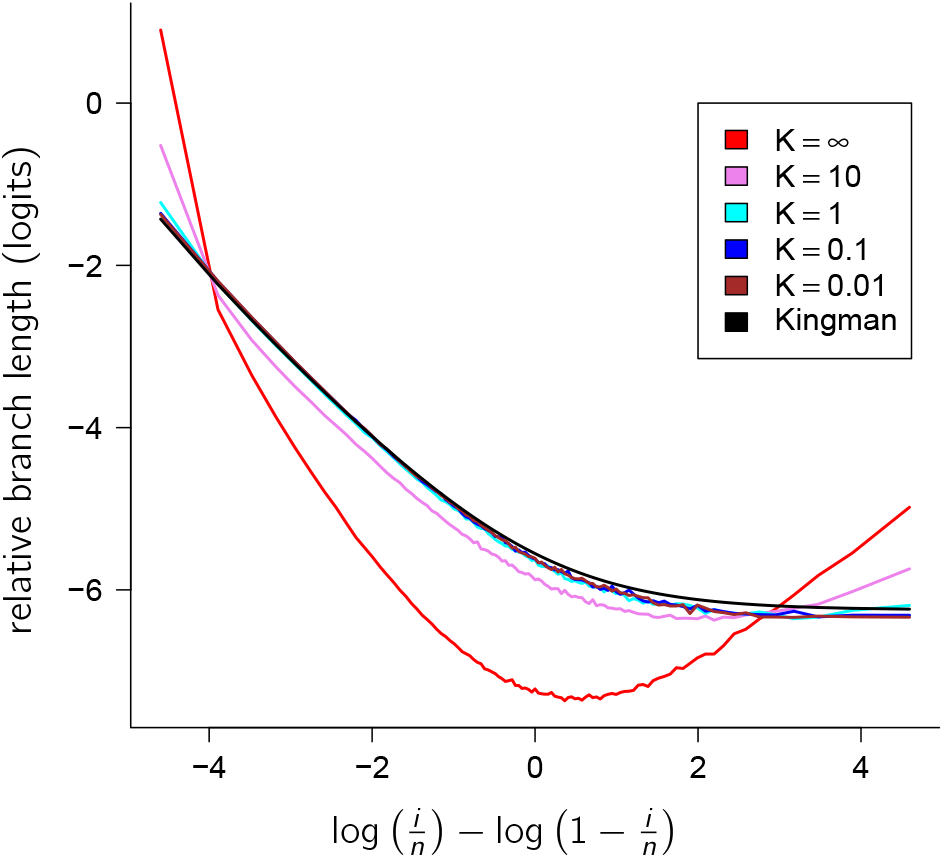
Comparing 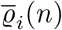 between Beta-coalescents. Estimates of 𝔼 [*R*_*i*_(*n*)] (recall (4.5) in Section 4) for sample size *n* = 100 predicted by the Beta(2 − *α, α*)-coalescent (*K* = ∞, red line), the incomplete Beta-coalescent for values of *K* (recall (3.12)) as shown and with *m*_∞_ approximated as in (4.3), and the Kingman coalescent (black line). For all the Beta coalescents *α* = 1.01; results from 10^5^ experiments.

## 5 Conclusion

Our main results are *(i)* the extension (3.2) of (1.1); our extension or some variant of it (see e.g. Remark 3.5) should be applicable to a broad class of populations. In particular, when *α ≥* 2 or *ψ*(*N*)*/N* → 0 the ancestral process is in the domain of attraction of the Kingman coalescent. *(ii)* Restricting to 1 *< α <* 2, then depending on the upper bound *ψ*(*N*) the limiting coalescent is a Kingman coalescent (Proposition 3.3), an incomplete Beta(2 − *α, α*) coalescent, or the complete Beta(2 − *α, α*) coalescent (Proposition 3.4); *(iii)* The error in the coalescent approximation can be quite large, especially if *ψ*(*N*) has a log *N* term (e.g. (3.10) and (3.11)); *(iv)* the effective size (recall Definition 2.5) can be much smaller than the population size with the ancestral process nevertheless in the domain of attraction of the Kingman coalescent (Case 1 of Proposition 3.2), for example if *ψ*(*N*) is of the form *N/* log *N*; *(v)* the sample size at which the gene genealogies of the ancestral process start to deviate from the Kingman trees can be much smaller than for the Wright-Fisher model, especially if the upper bound is of a specific form (see (3.17)); *(vi)* when the number of potential offspring are not bounded and the ancestral process is in the domain of attraction of the complete Beta-coalescent one can expect deviations in the topology of gene genealogies from the one predicted by the complete Beta(2 − *α, α*)-coalescent when sample size is at least *cN* ^*α/*2^ (some *c* > 0 fixed; see (8.5)). site-frequency spectrum depends on the relation between the sample size and the effective population size when the model is in the domain of attraction of the Kingman coalescent (Figure 3); increasing sample size does not distort the site-frequency spectrum when the model is in the domain of attraction of a Beta-coalescent (Figure 4). Thus, when the sample size is large enough that the gene genealogy should deviate from the limiting coalescent tree (e.g. Corollary 3.7, and the *cN* ^*α/*2^ result from Section 8.1) the site-frequency spectrum appears not to be affected. *(viii)* Applying the wrong Beta-coalescent in inference can lead to misinference (Figure 5 in Appendix C). *(ix)* Conditioning on the population ancestry does have an effect on the site-frequency spectrum for the models considered here (Figure 6 in Appendix D).

**Figure 6:**
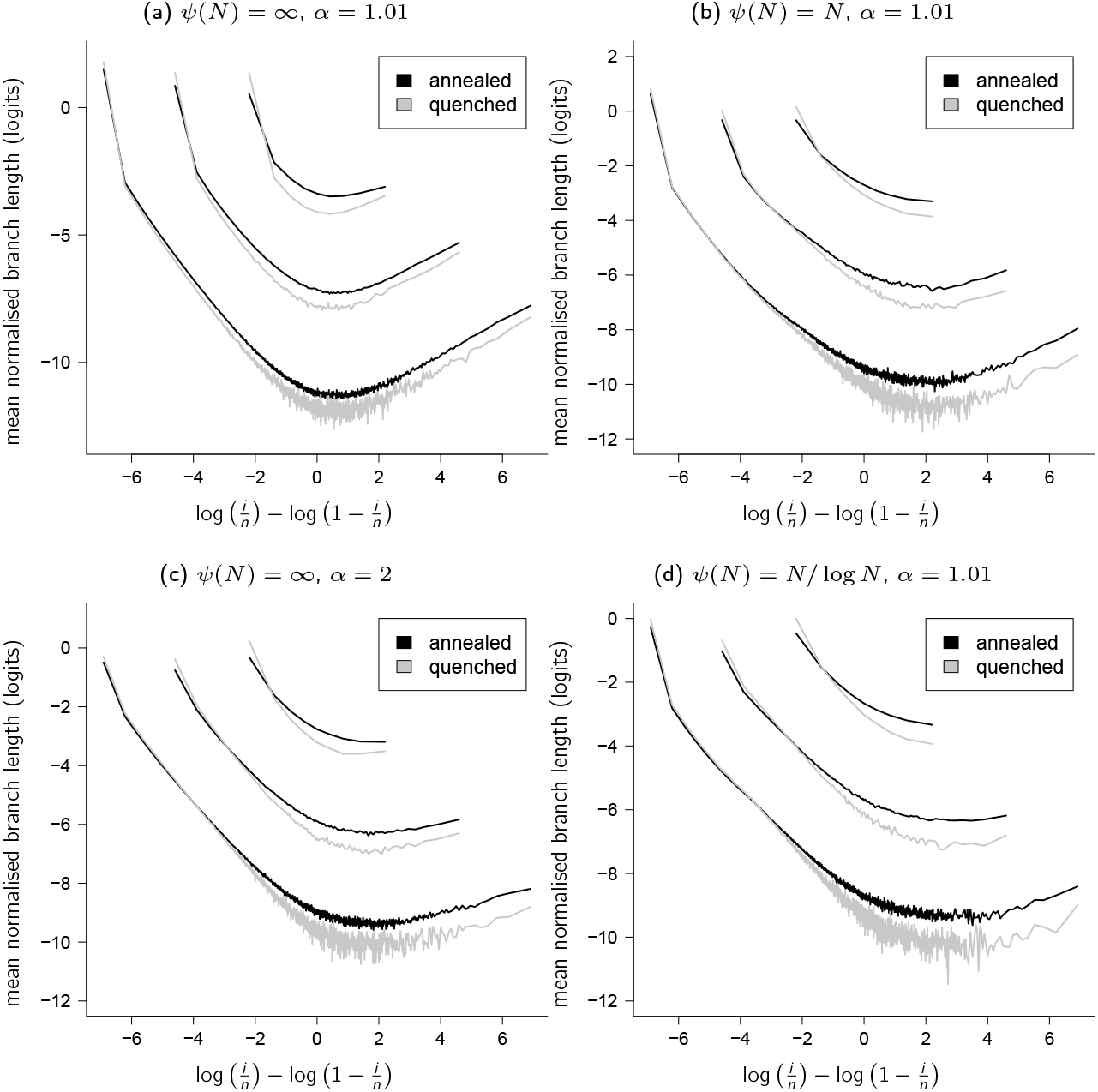
Comparing 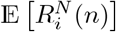 and 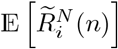. Estimates of mean relative branch lengths for *N* = 10^3^, *α* and *ψ*(*N*) as shown, sample size *n* = 10^1^, 10^2^, 10^3^; graphing 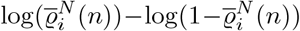 as a function of log(*i/n*) − log(1 − *i/n*) for *i* ∈ [*n* − 1] where 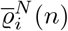 is an estimate of 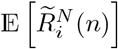 (quenched; grey) and 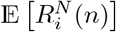 (annealed; black); results from 10^4^ resp. 10^5^ experiments. The limiting coalescent of the unconditional ancestral process (predicting 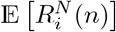) would be the complete Beta coalescent (a), incomplete Beta coalescent (b), and the Kingman coalescent (c,d). An algorithm for approximating 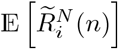 is briefly described in Appendix F and for 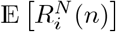 in Appendix E

In our model, a potential offspring is seen not as a gamete, but (at least) as a fertilised egg, and for many populations it would seem plausible that the number of fertilised eggs would, on average, be quite smaller than the number of gametes. By extending (1.1) to include an upper bound on the number of potential offspring we incorporate an important ecological reality, especially of broadcast spawners. Restricting *α* to (1, 2) leaves it to the upper bound to determine the limiting coalescent.

The coalescents we have considered are obtained by ignoring the ancestry, the ancestral relations, of a given sample. At the time of sampling (and all previous times) the ancestry of the entire population is fixed, but we derive our coalescents by averaging over the ancestry. Gene genealogies when the pedigree (ancestral relations between diploid individuals) is fixed (“quenched” or conditional gene trees) have been considered in diploid biparental organisms evolving according to the Wright-Fisher model, where the predictions of the quenched trees are shown to be similar to the ones of the classical (or “annealed”) Kingman coalescent (Wakeley et al., 2012). However, when the underlying (diploid) population evolves according to a particular model of sweepstakes reproduction, Diamantidis et al. (2024) show that the coalescent time of two gene copies, when conditioning on the population pedigree, does not converge to the time predicted by the corresponding multiple-merger coalescent. It is, therefore, of interest to investigate conditional gene genealogies in populations with sweepstakes reproduction. Here we consider haploid populations. In Appendix D we investigate if predictions of genetic variation will be different when conditioning on the population ancestry, even when the population is haploid. Figure 6 records an example comparing approximations of 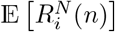 and 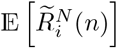, where we use 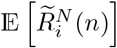 to denote the mean relative branch lengths read off fixed complete trees (see Appendix F for a brief description of the algorithm for approximating 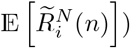. Even though the approximations broadly agree for the models considered in Figure 6, there is a noticeable difference. A quenched multiple-merger coalescent would be required to properly investigate the effect of increasing sample size on the predictions of conditional gene genealogies, and this we leave to future work, as well as the task (Árnason et al., 2023; Eldon, 2020; Freund et al., 2023) of extending our model to include diploidy (Möhle and Sagitov, 2003; Birkner et al., 2018, 2013a), many chromosomes, and complex demography in the spirit of Koskela and Berenguer (2019) and Freund (2020).

## 6 Key Lemmas

Throughout, the convergence (as *N* → ∞) of the pre-limiting (time rescaled) ancestral process {*ξ*^*N,n*^(⌊*t/c*_*N*_⌋); *t* ≥0} to a coalescent {*ξ*^*n*^} is in the sense of convergence of finite-dimensional distributions.

First we recall key lemmas. Our model is easily recast as a Cannings model (Cannings, 1974, 1975), enabling us to make use of Möhle and Sagitov (2001). For completeness, but following Schweinsberg (2003), we record the results that we need here. First we fix notation by describing the Cannings model for reproduction in a haploid panmictic population of constant size *N* evolving in discrete generations (Cannings, 1974, 1975). Write *ν*_*k*_ for the number of *surviving* offspring of the *k*th individual in an arbitrary generation, <u>*ν*</u> ≡ (*ν*_1_, *ν*_2_, …, *ν*_*N*_) is an exchangeable random vector with 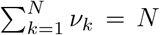. The vectors <u>*ν*</u> are assumed to be independent and identically distributed (abbreviated i.i.d.) across generations. Throughout we will be considering a haploid panmictic population of constant size *N* evolving as in Definition 2.3 and (3.2) (recall Remark 3.1).

Recall *c*_*N*_ from Definition 2.4. We see that for the Cannings model

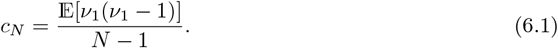

We now record general criteria for the convergence of the corresponding rescaled ancestral processes. The first, due to Möhle (2000), guarantees convergence to the Kingman coalescent. Recall the standard notation for descending factorials in (2.3) in Definition 2.1.

### Proposition 6.1

(Möhle (2000), Section 4). *Suppose, recalling c*_*N*_ *from* (6.1),

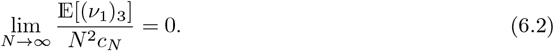

*Then c*_*N*_ → 0, *and for each finite sample size n* ∈ {2, 3, …} *the rescaled ancestral processes* {*ξ*^*n,N*^ (⌊*t/c*_*N*_ ⌋), *t* ≥ 0 *converge as N* → ∞ *to the Kingman coalescent restricted to* {1, …, *n*}.

If the family-size distribution is such that on the timescale determined by *c*_*N*_ we see ‘large families’ (comprising a non-trivial proportion of the total population) then we recover a coalescent with multiple mergers. Proposition 6.2 follows from Theorem 3.1 of Sagitov (1999).

### Proposition 6.2

(Sagitov (1999)). *Suppose both*

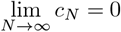

*and*

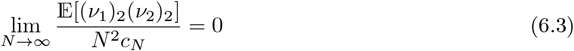

*hold. If there exists a probability measure* Λ_+_ *on* (0, 1] *such that for all x* ∈ (0, 1]

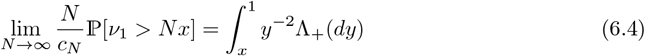

*then, for each finite sample size n* ∈ {2, 3, … }, *the rescaled ancestral processes* {*ξ*^*n,N*^ (⌊*t/c*_*N*_⌋), *t* ≥0} *converge as N* → ∞ *to a* Λ*-coalescent restricted to* {1, …, *n*} *admitting asynchronous mergers of a random number of lineages at such times*.

## 7 Asymptotic rate of the coalescent: calculating *c*_*N*_

First we set notation.

### Notation 7.1.

*Recall S*_*N*_ *from* (3.1). *As we are interested in the behaviour of*

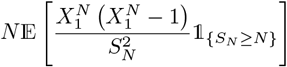

*it is convenient to fix a notation for* 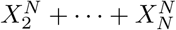; *we define*

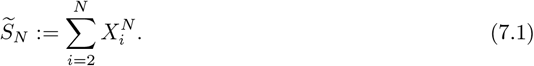

*Throughout* (*β*_*N*_)_*N*∈ℕ_ *will denote an increasing positive sequence. We will be partitioning over events (recall m*_*N*_ *from* (3.4))

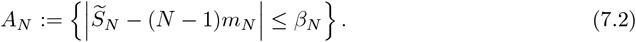

*and*

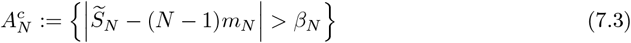

*(recall* 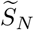 *from* (7.1)), 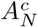 *being the complement of A*_*N*_. *Write*

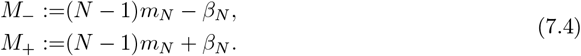

In this section we give a proof of Proposition 3.2. The quantity *c*_*N*_ is defined in Definition 2.4, recall also (6.1) and (3.9) for representations of *c*_*N*_.

### 7.1 Preliminary lemmas

#### Lemma 7.2.

*Suppose G and H are positive functions on* [1, ∞), *G be monotone decreasing and H monotone increasing, and that ∫ HG*′ *exists. Then*

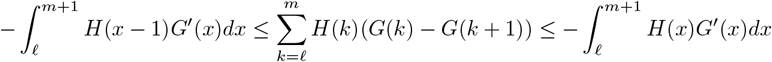

*Proof of Lemma 7.2*. It holds that

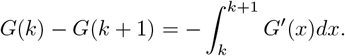

Then,

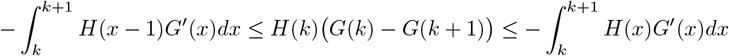

and all these quantities are non-negative. It remains to sum over *k* ∈ {*ℓ, ℓ* + 1, …, *m*}.

#### Lemma 7.3

(Bounding ∑_*k*_ (*k*)_2_(*k* + *M*)^−2^ (*k*^−*α*^ − (*k* + 1)^−*α*^)). *For all k* ∈ ℕ, *with M >* 1 *fixed, and* 1 *< α <* 2, *it holds that*

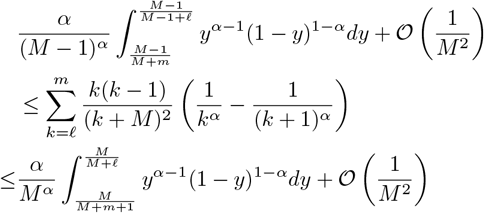

*Proof of* *Lemma* 7.3. Substituting *k*(*k* − 1)(*k* + *M*)^−2^ for *H* and *k*^−*α*^ for *G* in Lemma 7.2 gives

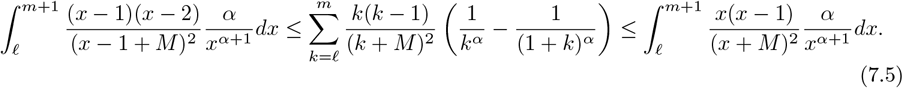

We evaluate the integrals appearing in (7.5). For the integral on the left in (7.5) we obtain, using the substitution *y* = (*M* − 1)*/*(*x* + *M* − 1),

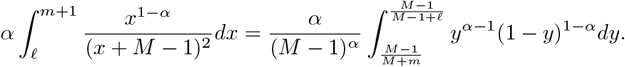

Integration by parts and the same substitution give us

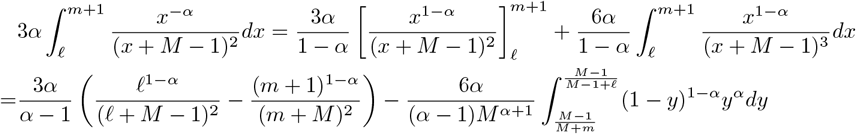

and using integration by parts twice and then the same substitution gives

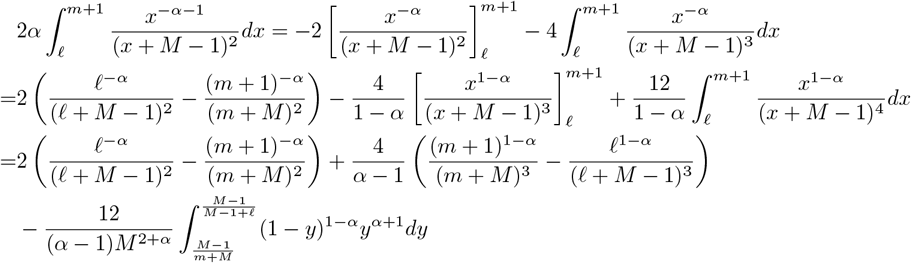

Similar calculations for the integral on the right in (7.5) give

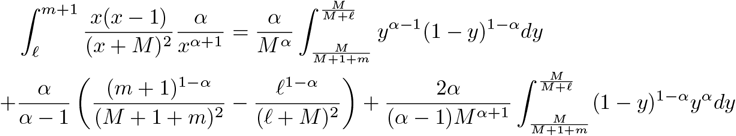

#### Lemma 7.4.

*For* 1 *< α <* 2, *any constant M >* 1, *with* 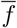 *and f defined in* (3.5), *and* 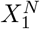 *distributed as in* (3.2), *and for any positive function L* ≡ *L*(*N*) ≤ *ψ*(*N*),

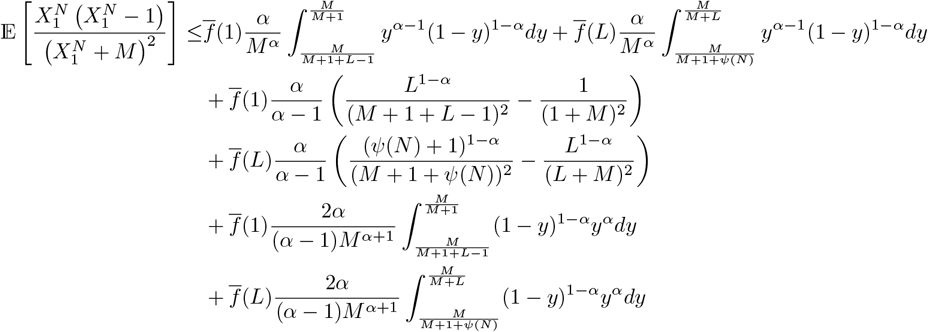

*and*

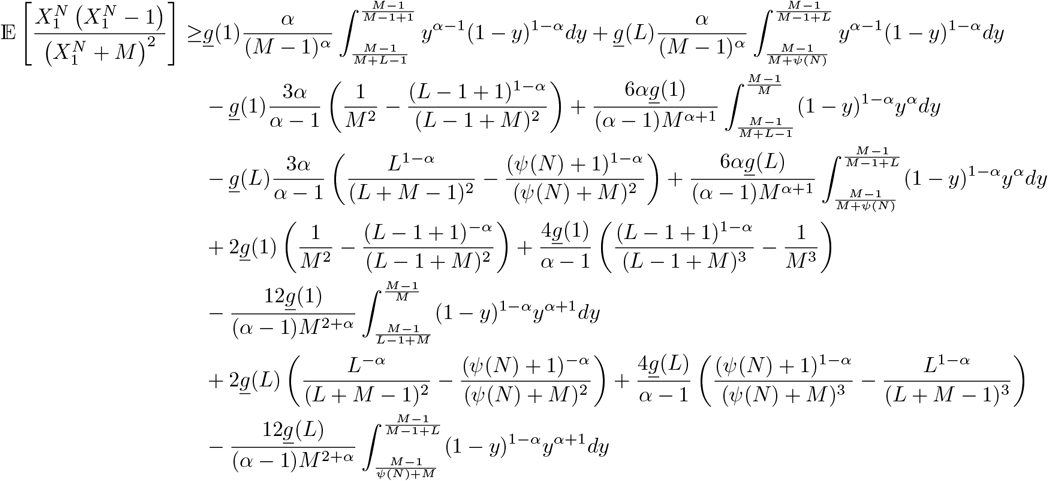

*Proof of Lemma 7.4*. The lemma follows from (3.2) and Lemma 7.3 and we recall 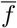 and *<u>f</u>* from (3.5); by (3.2)

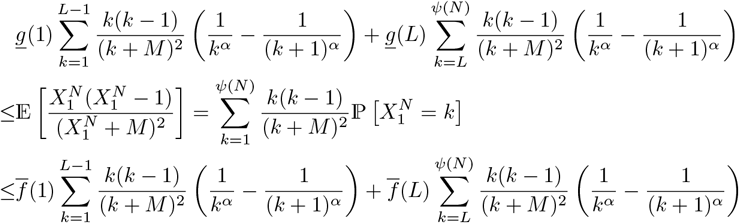

#### Remark 7.5

(The integrals in Lemma 7.4 and *ψ*(*N*)). *The integrals in Lemma 7.4 will all tend to zero unless ψ*(*N*)*/N* ≩ 0.

Proposition 3.2 will follow from Proposition 7.6 as an easy corollary. Section 7.2 contains a proof of Proposition 7.6.

#### Proposition 7.6.

*Suppose that the population evolves according to Definition 2.3 and* (3.2) *with* 1 *< α <* 2. *Let L be a function of N as specified in each case, recall m*_*N*_ *from* (3.4), 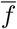 *and <u>f</u> from* (3.5) *and suppose f*_∞_ = *g*_∞_ *with f*_∞_, *g*_∞_ *from* (3.3). *Recall* 𝒞_*N*_ *from* (3.8).

1. *If ψ*(*N*)*/N* → 0 *and L*(*N*)*/ψ*(*N*) → 0 *then*

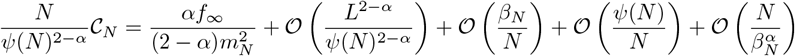
2. *If ψ*(*N*)*/N* → *K where K >* 0 *a constant, and L*(*N*)*/N* → 0 *then*

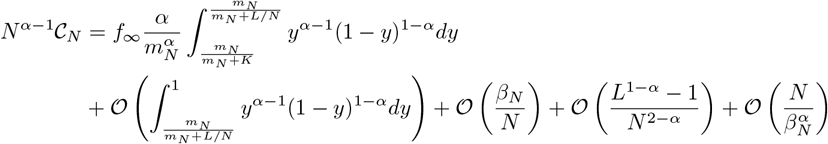
3. *If ψ*(*N*)*/N* → ∞ *and L*(*N*)*/N* → 0 *then*

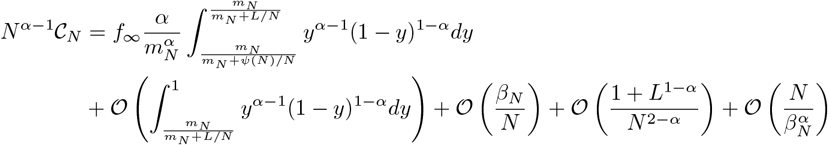

#### Remark 7.7.

*The constant order term in Case 3 reduces to a familiar form, since*

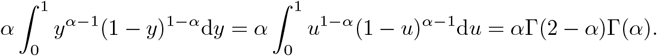

#### Remark 7.8

(The integral remainder term in Proposition 7.6). *Since L/N* → 0 *it holds that*

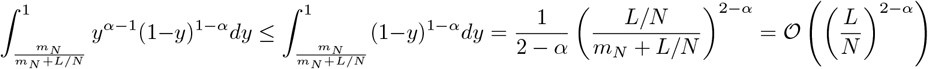

*as N* → ∞. *Moreover*,

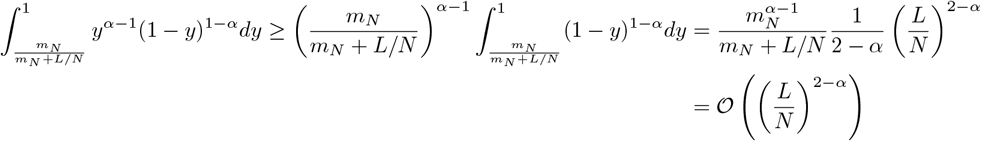

*as N* → ∞.

*Proof of Proposition 3.2*. This now follows immediately from Proposition 7.6 on setting

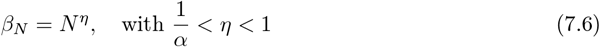

#### Remark 7.9

(*η* in (7.6)). *Examples of η in* (7.6) *such that* 1*/α < η <* 1 *include*

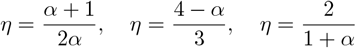

To prove Proposition 7.6 we will partition over the event *A*_*N*_ (defined in Notation 7.1), and its complement 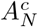. The random variable 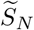 is defined in (7.1) in Notation 7.1; and *S*_*N*_ in (3.1). Our main focus will be the value, for large *N*, of the term

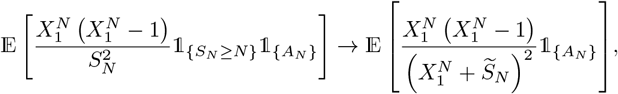

under the assumption that (recall *m*_*N*_ from (3.4)) *m*_*N*_ *>* 1 (so that ℙ [*S*_*N*_ *< N*] → 0 exponentially fast (Schweinsberg, 2003, Lemma 5)) and *β*_*N*_ */N* → 0. Proposition 7.6 will then follow if we prove that, recalling 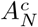 from (7.3),

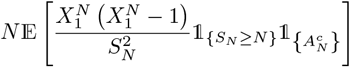

is asymptotically sufficiently small.

#### Lemma 7.10.

*In the notation of Proposition 7.6*, (7.3) *in Notation 7.1, as N* → ∞

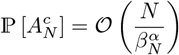

#### Remark 7.11.

*Taking ψ*(*N*) = ∞ *the rate in Lemma 7.10 is optimal as* 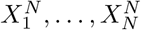 *will then lie in the domain of attraction of a stable law of index α for* 0 *< α <* 2 *and so we expect*

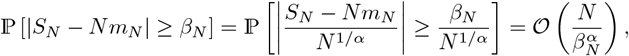

*for large N (Hall, 1981; Phillips, 1985)*.

*Proof of Lemma 7.10*. Define

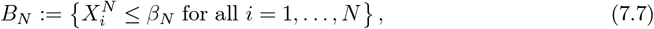

and observe that, recalling 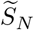 from Notation 7.1, and with 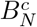 the complement of *B*_*N*_,

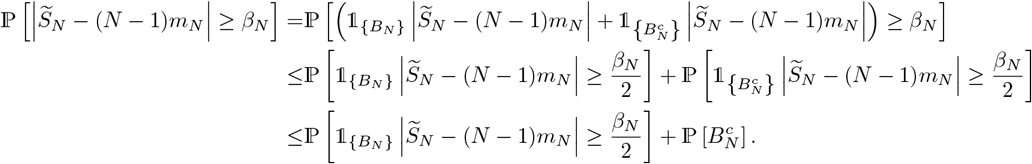

We can see from (3.2) and (3.6),

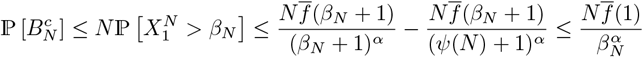

Define 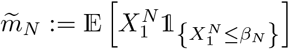. Then, with *m*_*N*_ from (3.4) and 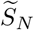 from (7.1)

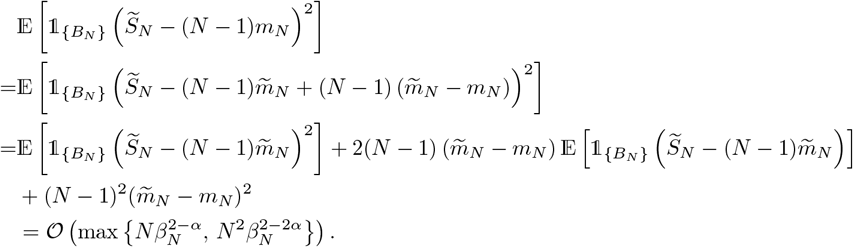

To explain the last line above we have by independence of the 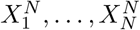

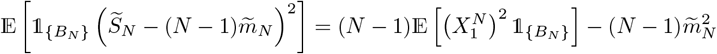

Using (3.2) we have (recall *B*_*N*_ from (7.7), and that (*β*_*N*_) is an increasing positive sequence)

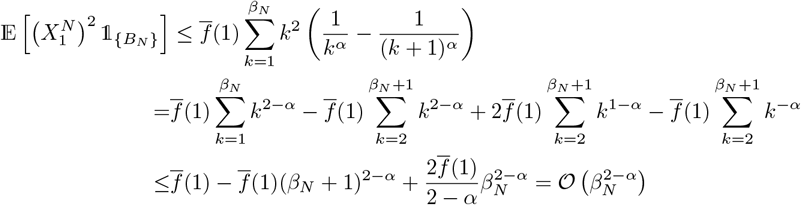

Similarly we have 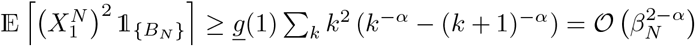 so that 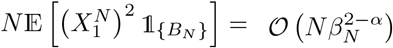. Since *α >* 1 we have lim 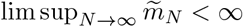 so that 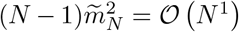. It follows that 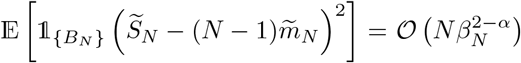. Since 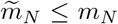 the term 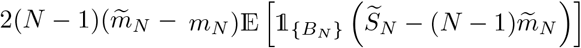 can be discarded; and 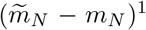 can be bounded by a constant times 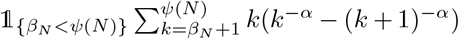, which is of order 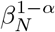 (use Lemma 7.2 with *H*(*x*) = *x* and *G*(*x*) = *x*^−*α*^).

Chebyshev’s inequality then implies

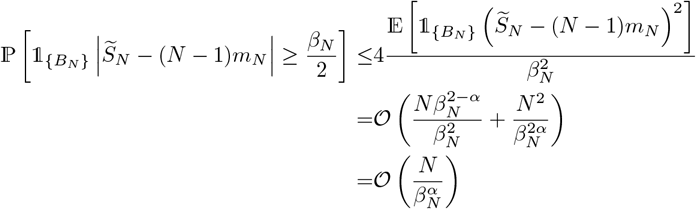

For 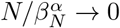. This concludes the proof of Lemma 7.10.

### 7.2 Proof of Proposition 7.6

We are now in a position to prove Proposition 7.6.

*Proof of Case 1 of Proposition 7.6*. We see using (3.2)

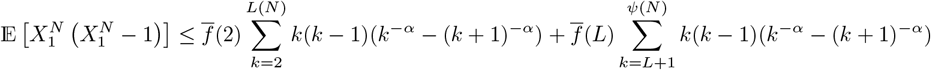

For *ℓ, m* ∈ ℕ with *ℓ* ≤ *m*

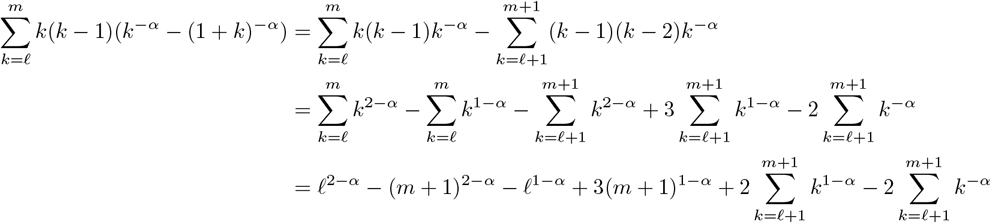

Since 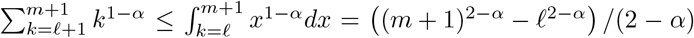 we have upon substituting for *ℓ* and *m* and writing *L* for *L*(*N*)

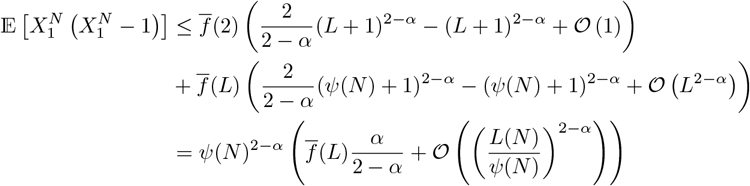

Similar calculations give

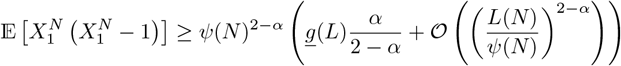

Recall *A*_*N*_ from (7.2) and *m*_*N*_ from (3.4) and *M*_−_ from (7.4) and 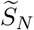 from (7.1). On *A*_*N*_ we have 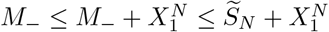. Choose *N* large enough that *S*_*N*_ ≥ *N* on *A*_*N*_. Then

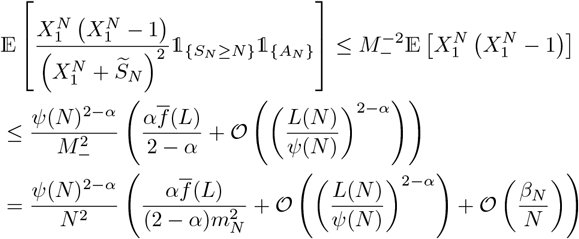

With *M*_+_ from (7.4) we have 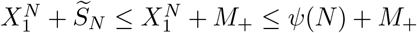 on *A*_*N*_. Then choosing *N* as before we have

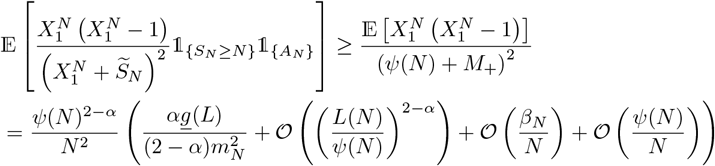

Recalling 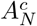 from (7.3), and that 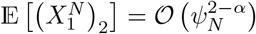 by our calculations above,

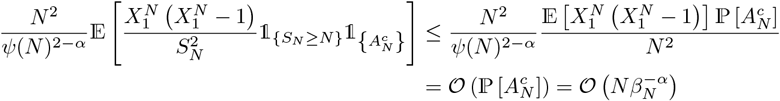

by Lemma 7.10. The proof of Case 1 of Proposition 7.6 is complete.

*Proof of Case 2 of Proposition 7.6*. Recall the assumption *ψ/N* → *K* with *K >* 0 constant, and the notation *M*_−_ from (7.4) in Notation 7.1, and *f*_∞_ from (3.3). We use Lemma 7.4 to see that if *L*(*N*)*/N* is bounded, and that 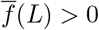 for any *L* by assumption and recalling that by (7.6) we have *β*_*N*_ */N* → 0 (and assuming *N* is large enough that *N* − 1 ≈ N)

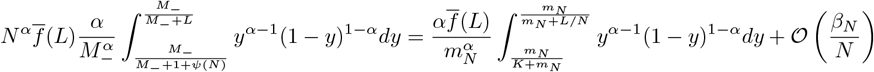

We then see

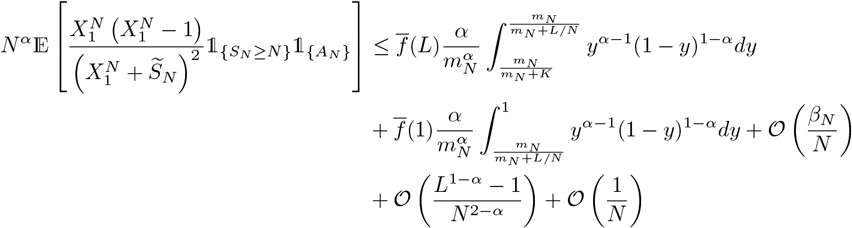

We also see, recalling *M*_+_ from (7.4),

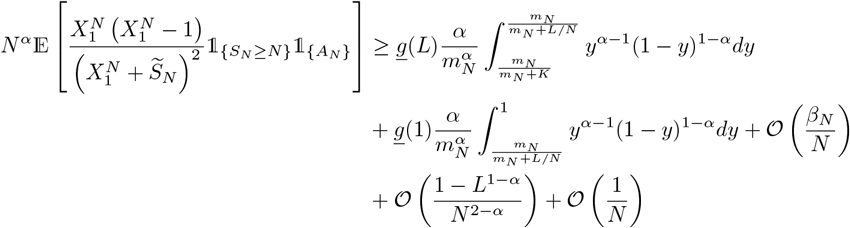

It remains to check that

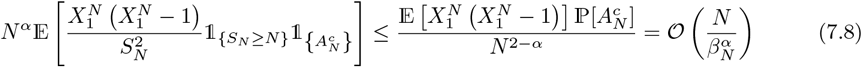

by Lemma 7.10 and the assumption *ψ*(*N*)*/N → K*. The proof of Case 2 of Proposition 7.6 is complete.

*Proof of Proposition 7.6, Case 3*. Now assuming that *ψ*(*N*)*/N* → ∞ and *L*(*N*)*/N* bounded, we again turn to Lemma 7.4 to see

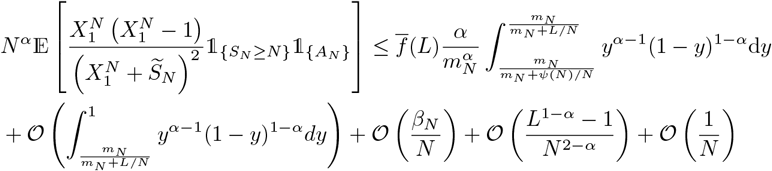

Similarly,

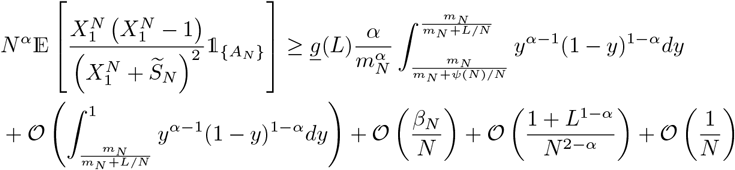

When *S*_N_ ≥ *N* then 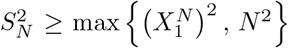, and 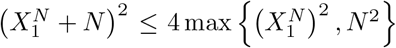 always holds. With this in mind we see

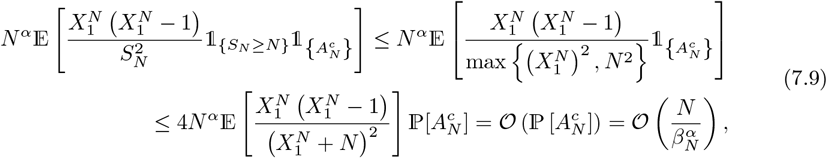

where we have used Lemma 7.10 to bound 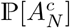 and Lemma 7.4 to control all other terms. The proof of Case 3 of Proposition 7.6 is complete.

We conclude this section with a slight modification of Lemma 6 in (Schweinsberg, 2003), and a proof of (3.9).

#### Lemma 7.12

(Schweinsberg (2003), Lemma 6). *Suppose a haploid population of size N evolves according to Definition 2.3. Recall our notation ν*_1,*N*_, …, *ν*_*N,N*_ *for the random number of surviving offspring of the N parents, in a given generation, where ν*_1,*N*_ + · · · + *ν*_*N,N*_ = *N. For all r* ≥ 1 *and k*_1_, …, *k*_*r*_ ≥ 2, *we have*

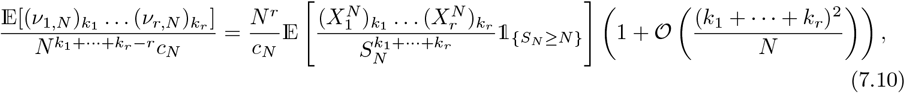

*where c*_*N*_ *is defined in Definition 2.4*, 𝒞_*N*_ *in* (3.8), *and*

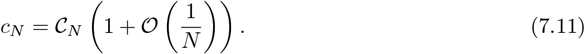

Our proof follows that presented in Schweinsberg (2003). However, we include it in order to emphasize that the source of the error term in (7.10) is the distinction between sampling with and without replacement from the juvenile population.

*Proof of Lemma 7.12*. Label individuals in the current generation by {1, 2, …, *N* }. For each 1 ≤ *r* ≤ *N*, set *k*_0_ = 0 and write 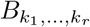 for the event that for each *i* = 1, …, *r*, the individuals labelled {*k*_0_ + ···+ *k*_*i*−1_ + 1, …, *k*_0_ + ···+ *k*_*i*−1_ + *k*_*i*_ } are all descended from the individual with label *i* in the previous generation.

We then see that

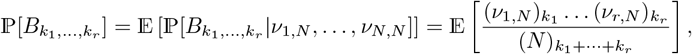

and

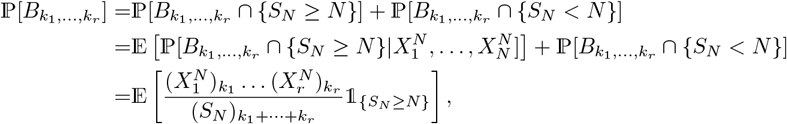

where we have used that when *S*_*N*_ *< N* we will keep the previous population, and so will see exactly one individual in the *i*th generation descended from each individual in the previous generation.

The result follows on replacing 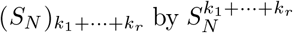; expanding the falling factorial (*n*)_*m*_ (see (2.3)) gives (recall 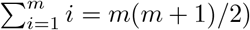

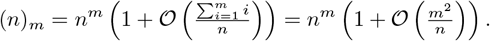

We recall the definition of 𝒞_*N*_ from (3.8). We may deduce (7.11) from (7.10) with *r* = 1 and *k*_1_ = 2, by using that *c*_*N*_ = 𝔼 [(*ν*_1,*N*_)_2_]*/*(*N* − 1) and that, in this case, the left hand side of (7.10) is equal to (*N* − 1)*/N*. This concludes the proof of Lemma 7.12.

## 8 Proof of Lemma 3.6

Recall the notation in Definition 7.1 in Section 7, *M*_−_ and *M*_+_ are defined in (7.4), 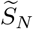 in (7.1) and *A*_*N*_ in (7.2).

We will use the notation *M*_−_ and *M*_+_ throughout our calculations. In Lemma 8.1 and in (8.2) we use *M* to denote an arbitrary (positive) constant. We then intend to use the result with *M* replaced by either *M*_+_ or *M*_−_.

For any natural numbers *N* ≥ *n* ≥ *k* ≥ 2 let *E* be the event

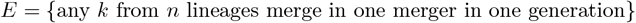

Then, conditioning on 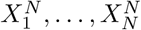,

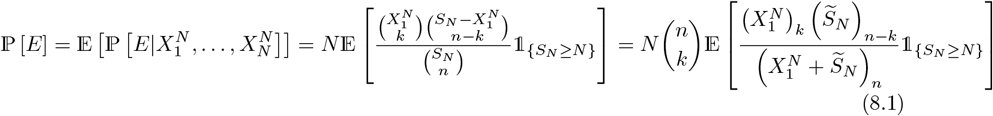

and we have not prohibited simultaneous mergers.

As usual we consider the set *A*_*N*_ defined in (7.2) in Notation 7.1. On *A*_*N*_, for any *n* with *n/S*_*N*_ → 0, we approximate 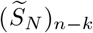 by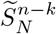, and 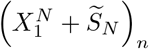 by 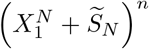.

We will make use of the following observation: for any 0 *< a < b <* ∞, *M >* 0, and using the substitution *y* = *M/*(*M* + *x*) (so that *x* = *M/y* − *M* and *dx* = −*My*^−2^*dy*),

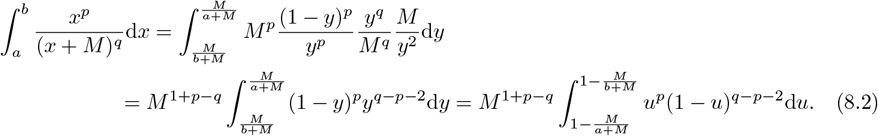

using the substitution *u* = 1 − *y* for the last equality. We also recall from the calculations in Section 10 that, for *ψ*(*N*)*/N* → ∞ and 0 *< x <* 1,

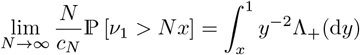

with Λ_+_(d*y*) = *C*(*α*)(1 − *y*)^*α*−1^*y*^1−*α*^d*y*, where the constant *C*(*α*) = 1*/*(Γ(*α*)Γ(2 − *α*)) (recall (6.4) in Proposition 6.2 in Section 6).

### Lemma 8.1.

*For M >* 0 *and* 0 *< a < b <* ∞ *constants we have, for p, q >* 0, *and p*≠ *q*,

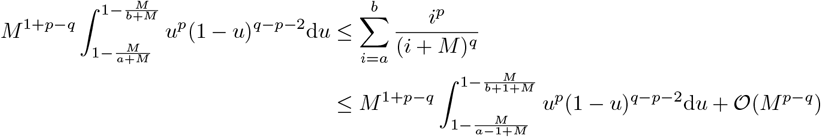

*Proof of Lemma 8.1*. We begin with, recalling (8.2),

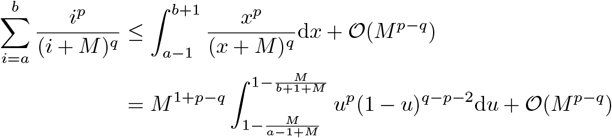

where the 𝒪 (*M* ^*p*−*q*^) correction appears since for *p*≠ *q, i*^*p*^*/*(*i* + *M*)^*q*^ has a maximum at *i* = *pM/*(*q* − *p*). A similar calculation provides the opposite inequality.

*Proof of Lemma 3.6*. As usual we partition over *A*_*N*_ (7.2) and 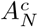 (7.3). On *A*_*N*_ it holds that

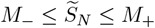

Recall (8.1), *M*_+_, *M*_−_ from (7.4), *S*_*N*_ from (3.1), 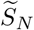 from (7.1), and note that

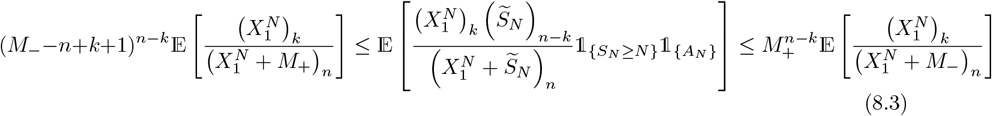

We will use that 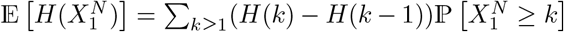 for a real-valued function *H H*(0) = 0 (Schweinsberg, 2003, Lemma 9), and the falling factorial (recall (2.3)) identities (*i*)_*k*_(*i* −*k*) = *i*(*i* − 1)_*k*_, (*i*)_*k*_ = *i*(*i* − 1)_*k*−1_, and (*i* − 1)_*k*_ = (*i* − 1)_*k*−1_(*i* − *k*). We now look at 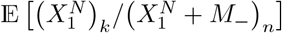 from (8.3). We see

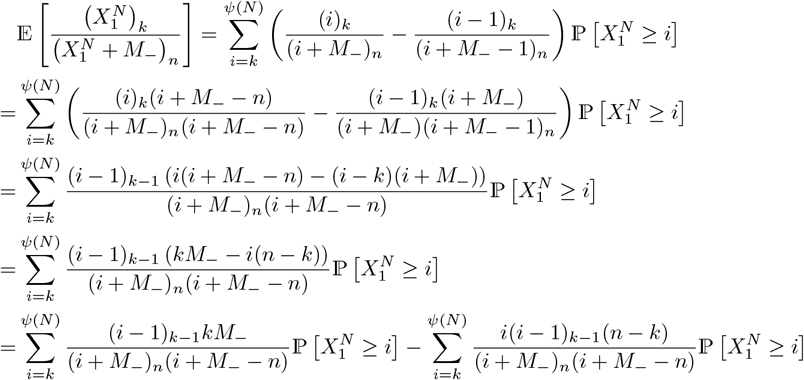

Our next task is to control the first term in this expression. For any *L* ≤ *ψ*(*N*) with *L/N* → 0,

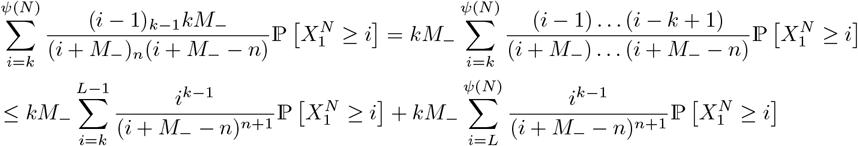

Lemma 8.1 and (3.6) now give (with 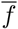, *f* from (3.5))

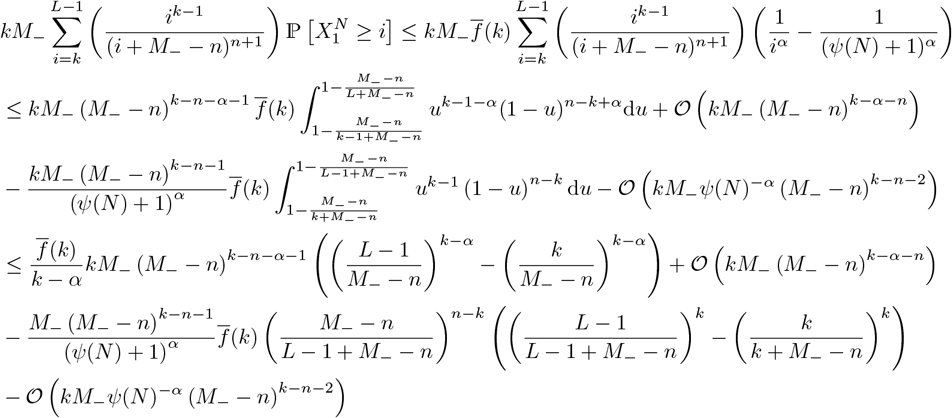

Again combining Lemma 8.1 and (3.6) gives

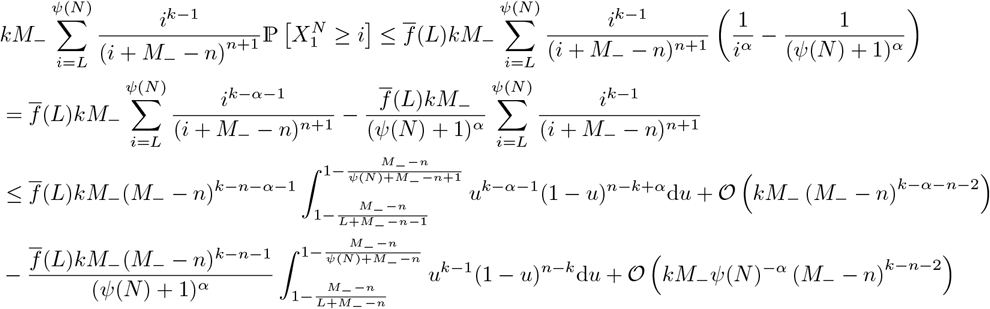

To find a corresponding lower bound, first observe that

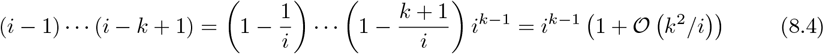

It then holds by (3.6) and Lemma 8.1

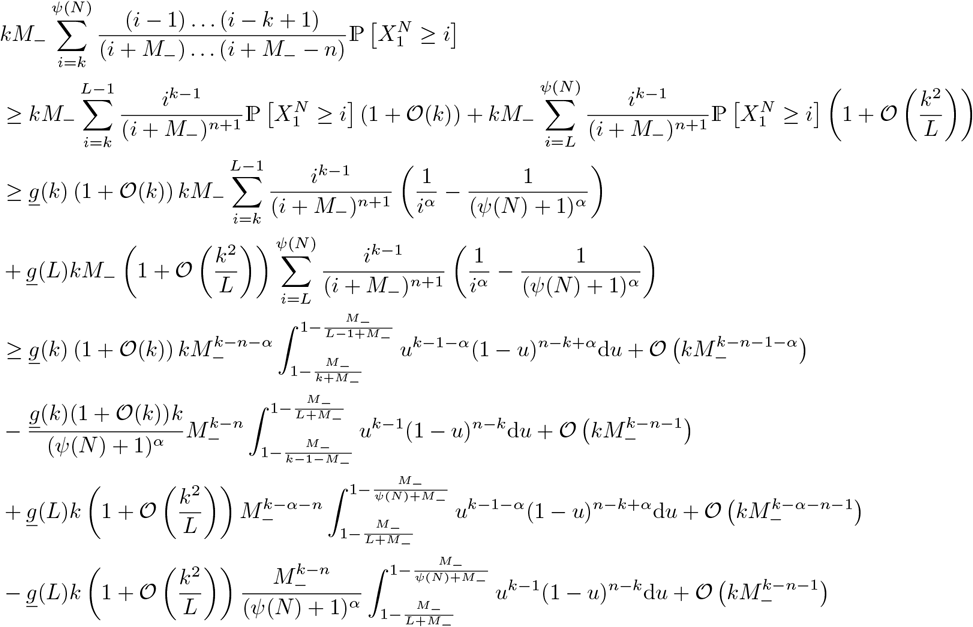

Analogous arguments yield

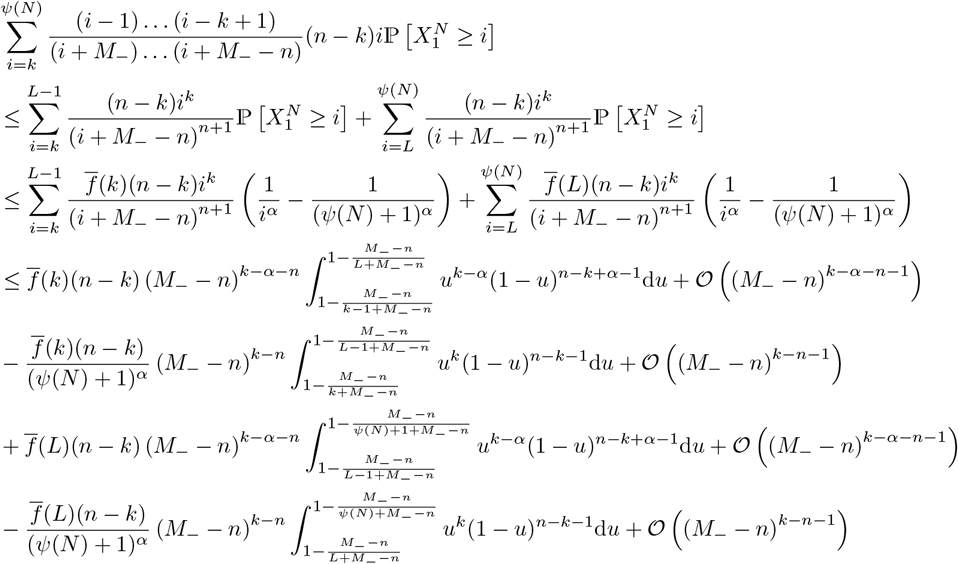

Similar calculations give us the lower bound (recall (8.4))

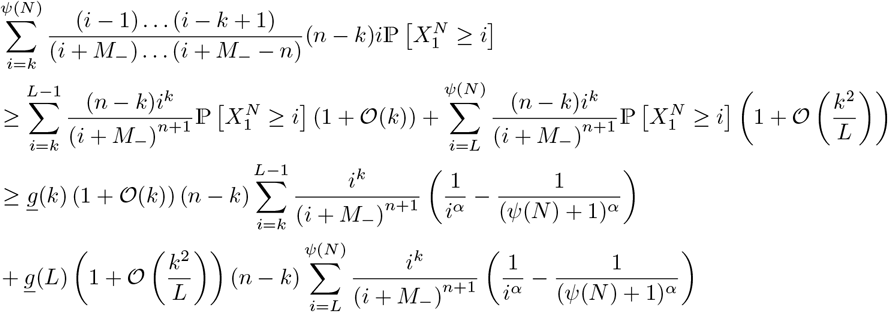

We can now apply Lemma 8.1 to get

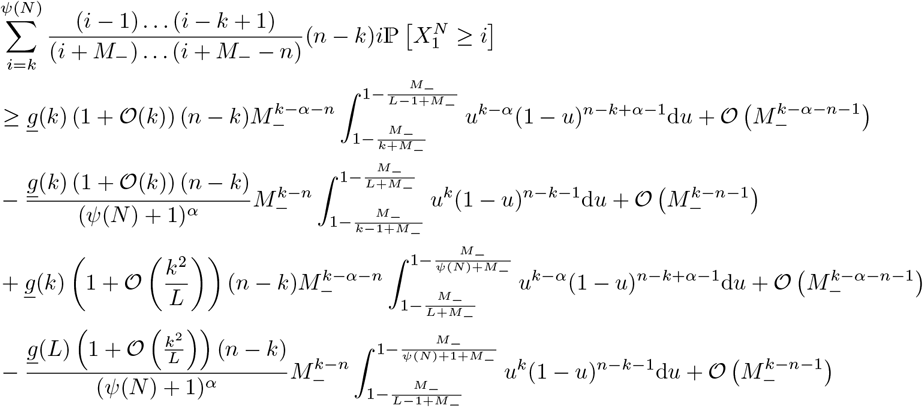

Bounds for 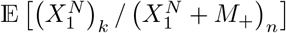 (recall (8.3)) follow from analogous arguments.

We must now estimate the terms corresponding to 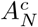 (7.3). When *n/N* → 0, for sufficiently large *N*, using the identity (*x*)_*n*_ = (*x*)_*n*−*k*_(*x* − *n* + 1) · · · (*x* − *n* + *k*),

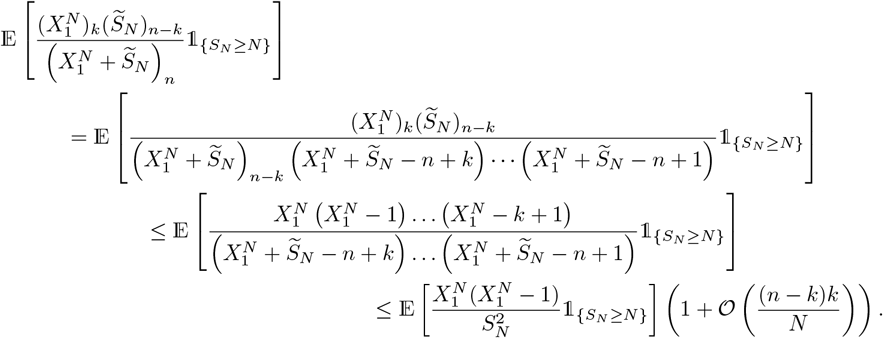

The error term is calculated by partitioning over 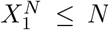 and 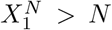. In the first case, each of the *k* ‘spare terms’ 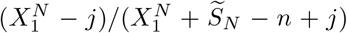 is bounded by *N/*(*N* − (*n* − *k*)); if 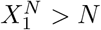 each such term is bounded by 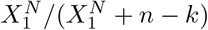. Now using the elementary facts that (*i* + *b*)^*k*^*/i*^*k*^ = 1 + 𝒪 (*kb/i*) and (*i* − *b*)^*k*^*/i*^*k*^ = 1 − 𝒪 (*kb/i*) we recover the error term above.

We now see that the error term 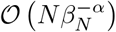 in (3.18) and (3.19) comes from (7.8) in the case *ψ*(*N*)*/N* → *K* and from (7.9) when *ψ*(*N*)*/N* → ∞. The proof of Lemma 3.6 is complete.

### 8.1 Heuristic arguments for sample size 𝒪 *N* ^*α/*2^

Here we give heuristics for the sample size when we would expect to start seeing anomalies in the topology of the tree predicted by the complete Beta(2 −*α, α*) coalescent (Definition 2.2).

Since

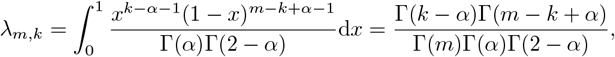

the total rate of a *k*-merger (with *k >* 2) when *m* lineages is given by

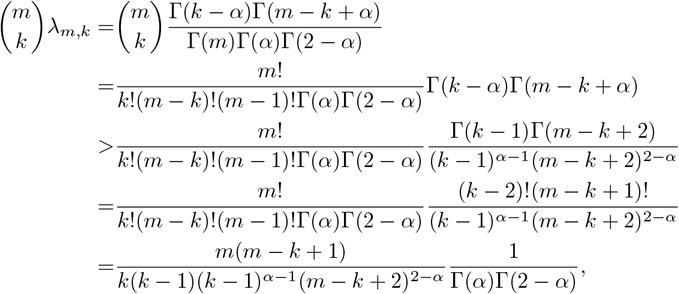

obtained by writing Γ(*k− α*) = Γ((*k−* 2) + (2 −*α*)) and Γ(*m− k* + *α*) = Γ((*m −k* + 1) + (*α −*1)) and using Gautschi’s inequality (Gautschi, 1959)

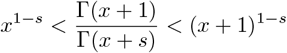

(so that Γ(*x* + *s*) *>*Γ(*x* + 1)*/*(*x* + 1)^1−*s*^) for *x >* 0 and *s* ∈ (0, 1).

Suppose 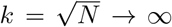. Approximating 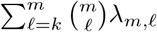 by an integral, the total rate of mergers involving at least *k* lineages can be approximated (up to a constant) by

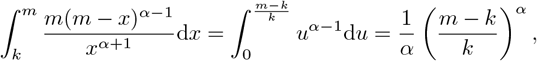

by the substitution *u* = (*m −x*)*/x* (so that *x* = *m/*(1 + *u*), and *dx* = −*m/*(1 + *u*)^2^*du*).

Now consider the number of ancestral lineages present in the Beta(2 − *α, α*) coalescent at any given time. We recall the following result from Berestycki et al. (2008b).

#### Theorem 8.2

(Berestycki et al. (2008b), Theorem 1.1). *Let* 𝔹_*t*_ *be the remaining number of blocks at time t in the complete Beta*(2 − *α, α*) *coalescent* (2.7) *with* 1 *< α <* 2. *Then*

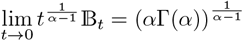

*holds almost surely*.

We now present some simple heuristic arguments. We will be looking at small time results and so, with an abuse of notation, we take (see Theorem 8.2)

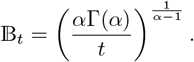

We then see that the expected number of mergers involving more than *k* individuals when started from *m* individuals is approximated by

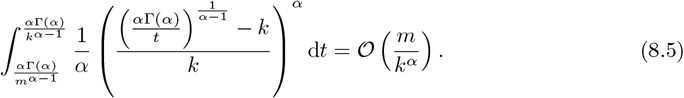

The integral is dominated by the contribution for times arbitrarily close to the starting time, so as soon as *m* = 𝒪 (*N* ^*α/*2^) we expect to see mergers of at least 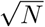 lineages at arbitrarily small times. Since the external branches of a Beta-coalescent extend into the ‘body’ of the tree, see e.g. (DAHMER et al., 2014) (unlike in the Kingman case in which the ratio of external to internal branches tends to zero), such mergers can be expected to influence the site-frequency spectrum for sample sizes of 𝒪 (*N* ^*α/*2^), as claimed. This holds even though 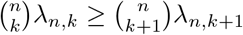 and

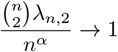

as *n → ∞*. In words, the merger probabilities are decreasing as a function of merger size, so a 2-merger is always the most likely merger, and the probability of seeing a 2-merger tends to one as sample size increases.

## 9 Convergence to the Kingman Coalescent

In this section we use Proposition 6.1, where the condition given in (6.2) requires us to control the probability of three ancestral lines merging, to prove Proposition 3.3.

*Proof of Proposition 3.3*. Recalling *S*_*N*_ from (3.1) and setting *β*_*N*_ = *N*, we see

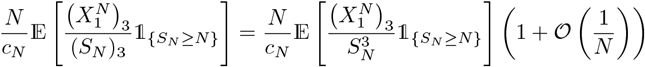

Then, according to Lemma 7.12, it will suffice to replace *c*_*N*_ by 𝒞_*N*_, and from Proposition 3.2 𝒞_*N*_ = 𝒪(*ψ(N)*^*2−α*^/*N*),. Recall that we are assuming that *ψ*(*N*)*/N* → 0. We then see

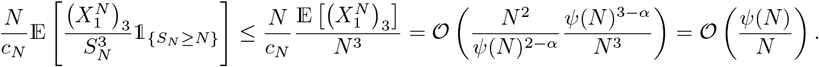

From this we can conclude convergence to Kingman coalescent by Proposition 6.1. To establish (3.10) it remains to check the lower bound. Recalling *A*_*N*_ from (7.2) and *M*_+_ from (7.4) and 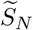 from (7.1), it holds that

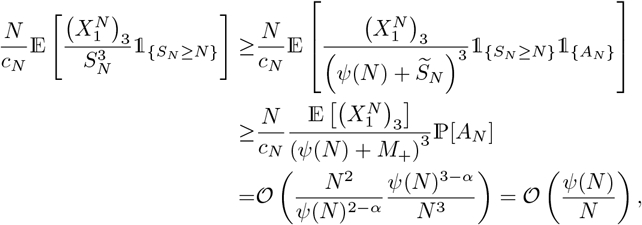

where we used Lemma 7.10 to deduce that ℙ [*A*_*N*_] is 𝒪 (1).

Considering (3.11), we obtain

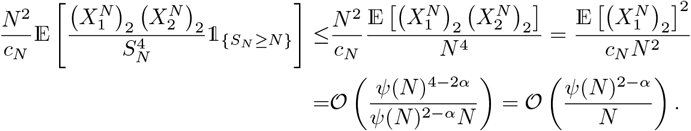

For the opposite inequality, we define

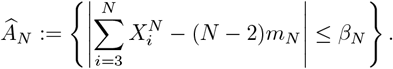

Proceeding as before we see

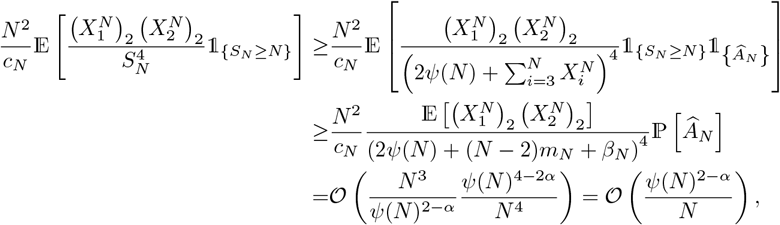

where again we chose *β*_*N*_ = *N*, and used Lemma 7.10. This concludes the proof of Proposition 3.3.

## 10 Convergence to a Beta-coalescent

In this section we use Proposition 6.2 to prove Proposition 3.4.

We divide the proof of Proposition 3.4 into two parts. First we control the probability of simultaneous multiple mergers (recall the condition in (6.3) in Proposition 6.2), and then we identify the measure Λ in the limiting coalescent, see (6.4) in Proposition 6.2.

*Proof of* (3.13). According to Lemma 7.12 we can replace *c*_*N*_ by 𝒞_*N*_ (see (3.8)), and according to Proposition 3.2 𝒞_*N*_ = 𝒪 (*N* ^1−*α*^) (recall 1 *< α <* 2). Then, assuming that *N* is large enough that *m*_*N*_ *>* 1 following Schweinsberg (2003) we see

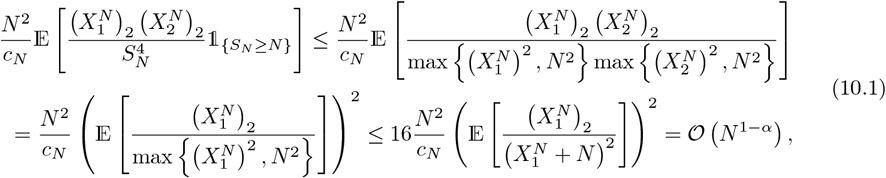

where we have used Lemma 7.4 for the final estimate.

We turn to the lower bound. Using the Notation 7.1 and *M*_+_ from (7.4) we have

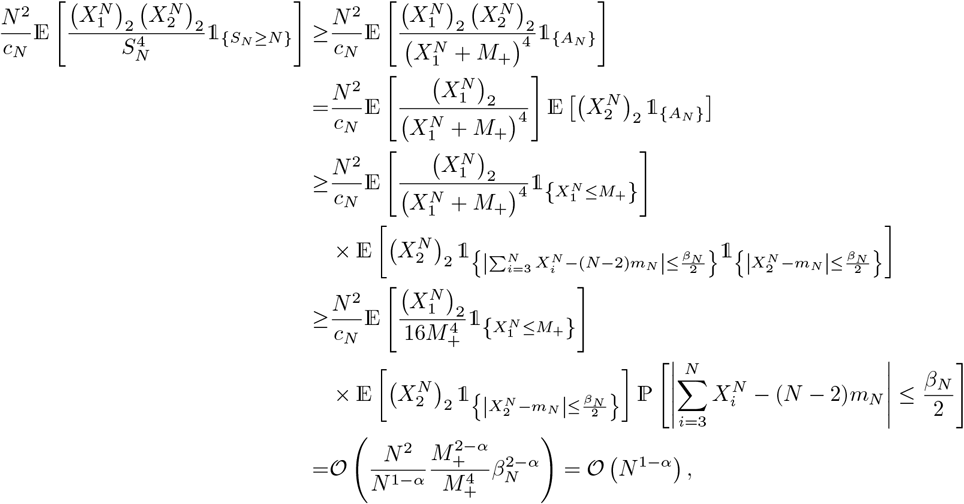

where we choose *β*_*N*_ = *N* to optimise on the final line and Lemma 7.10 to deduce that the probability on the penultimate line is 𝒪 (1). The proof of (3.13) is complete.

*Proof of Proposition 3.4: identifying the measure* Λ_+_. We first note that (3.13) and Lemma 7.12 reduce the proof of Proposition 3.4 to examining

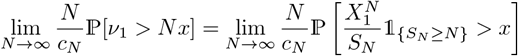

for 0 *< x <* 1. We essentially follow Schweinsberg (2003), pp 129–131. We start in a familiar way by conditioning on 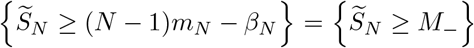. As before we assume *β*_*N*_ */N* is bounded (with *β*_*N*_ for example as in (7.6)), and recall the definition of *A*_*N*_ from Notation 7.1, and of *M*_+_ and *M*_−_ from (7.4). We see, using Lemma 7.10 for 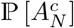, and that 1*/c*_*N*_ = 𝒪 (*N* ^*α*−1^) from Proposition 3.2,

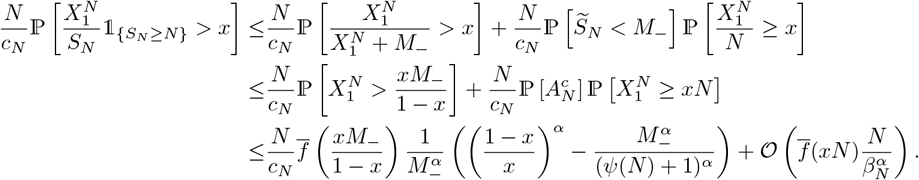

Considering a lower bound, we have

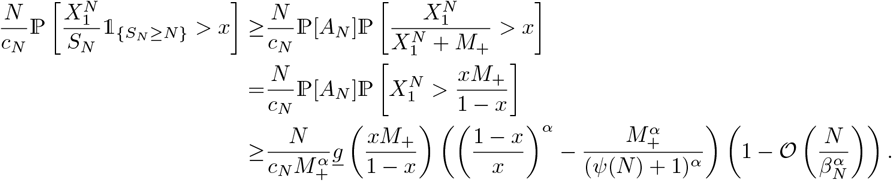

Combining these estimates with (10.1) we conclude from Proposition 6.2, assuming *β*_*N*_ */N* is bounded, convergence to the claimed coalescent; for any *A* ≥ 0, *r >* 0, and 0 *< x* ≤ 1*/*(1 + *A*),

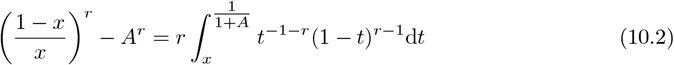

seen using the substitution *u* = (1 − *t*)*/t*. Since ∫*y*^−1−*α*^(1 *y*)^*α*−1^d*y* = ∫*y*^−2^ *y*^1−*α*^(1 *y*)^*α*−1^ d*y* we have, as required by Proposition 6.2, identified a measure Λ_+_ so that the limit in (6.4) holds. Using Lemma 7.12 to replace *c*_*N*_ by _*N*_ from Proposition 3.2, the proof of Proposition 3.4 is complete.

## Acknowledgements

JACD supported by an Engineering and Physical Sciences Research Council (EPSRC) grant number EP/L015811/1; BE supported in part by EPSRC grant EP/G052026/1 to AME, by Deutsche Forschungsgemeinschaft (DFG) grant BL 1105/3-1 to Jochen Blath through SPP Priority Programme 1590 Probabilistic Structures in Evolution, by DFG grant with Projektnummer 273887127 through DFG SPP 1819: Rapid Evolutionary Adaptation grant STE 325/17 to Wolfgang Stephan; AME and BE acknowledge funding by Icelandic Centre of Research (Rannís) through an Icelandic Research Fund (Rannsóknasjóður) Grant of Excellence no. 185151-051 jointly with Einar Árnason, Katrín Halldórsdóttir, Alison Etheridge, and Wolfgang Stephan; BE also acknowledges SPP 1819 Start-up module grants jointly with Jere Koskela and Maite Wilke Berenguer, and with Iulia Dahmer. We thank Alison M. Etheridge for helpful comments and suggestions, and especially for suggesting Lemma 7.2.

## Statements and Declarations

The authors have no competing interests to declare that are relevant to the content of this article.

## MSC Classification

60J10, 60J90, 60J95, 60-04

## Data availability

The software developed during this study is available at https://github.org/eldonb/Beta_coalescents_when_sample_size_is_large

## A Relative length of external branches in the Wakeley-Takahashi model and increasing sample size

In this section we provide further arguments for why one would see an increase in relative length of external branches in the Wakeley-Takahashi model (recall Section 3.3.1) with increasing sample size.

The following lemma and proof are a straightforward adaptation of the corresponding mean passage time bound derived by Ross and Peköz (2007).

### Lemma A.1.

*Consider a discrete-time Markov chain taking values in* ℕ *with transition probability matrix* (*p*_*i,j*_)_*i,j*_ *with p*_*i,j*_ = 0 *for i < j and j* = 0. *Let S*_*i*_ *denote the random sum of the states visited when started from state i* ≥ 2 *until state 1 is reached. Let* (*d*_*i*_)_*i*_ *be a strictly positive sequence with d*_*i*_ ≤ *d*_*i*+1_ *and i* ≤ 𝔼 [*D*_*i*_] */d*_*i*_ *for all i* ∈ ℕ, *where* ℙ [*D*_*i*_ = *k*] = *p*_*i,i*−*k*_, 0 ≤ *k* ≤ *i, i*.*e. D*_*i*_ *is the amount of decrease as the Markov chain transits from state i to state i* − *k. Then*

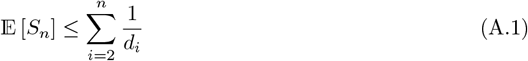

*Proof*. The proof proceeds by induction. We have 𝔼 [*S*_1_] = 0, and 𝔼 [*S*_2_] = 2 + 𝔼 [*S*_2_] ℙ [*D*_2_ = 0]. Since 1 − ℙ [*D*_2_ = 0] = ℙ [*D*_2_ = 1] = 𝔼 [*D*_2_] we have

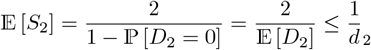

Assuming 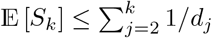 for all *k* ∈ {2, …, *n* − 1}, we see

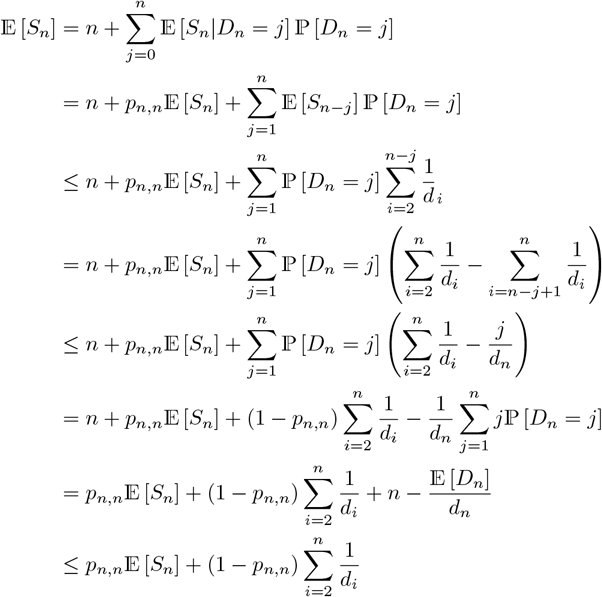

since *n* ≤ 𝔼 [*D*_*n*_] */d*_*n*_ by assumption.

The result in Theorem A.2 may be well known, but for lack of a point reference we include it here. In the context of (A.2) recall the well-known result 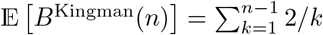, where *B*^Kingman^(*n*) is the random total tree size of a sample of size *n* under the Kingman coalescent (clearly lim sup_*n*→∞_ 𝔼 *B*^Kingman^(*n*) */* log *n <* ∞).

### Theorem A.2.

*Let B*^WF^(*n, N*) *denote the random tree size of a sample of size n* ≥ 2 *from a haploid panmictic population of constant size N evolving according to the Wright-Fisher model with time measured in generations. Then, with B*^WF^(*N, N*) *denoting the total tree length of the entire population when it evolves according to the Wright-Fisher model*,

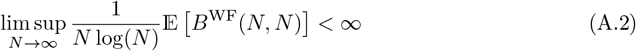

*Proof*. Consider a the Markov sequence representing the block-counting process of a sample of size *n* from a haploid population of constant size evolving according to the Wright-Fisher model. Let *S*_*n*_ represent the size of the tree of a sample of size *n*. Now we consider 𝔼 [*S*_*n*_] when *n* = *N*, where *N* is the population size. Following Ross and Peköz (2007) we have 𝔼 [*D*_*i*_] = *i* − *N* + *N* (1 − 1*/N*)^*i*^. Write *p*_*N*_ := 1 − 1*/N*. We see, for any *i* ∈ {2, …, *N* − 1},

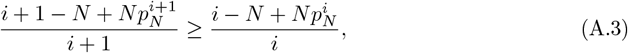

since, by rearranging and canceling terms, Eq (A.3) simplifies to

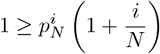

Eq (A.3) can be verified by induction. For *i* = 2 Eq (A.3) holds for all *N* ≥ 2*/*3. Assuming Eq (A.3) holds for a given *i* we see

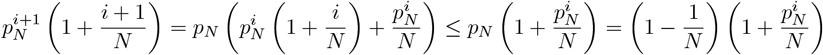

where the inequality follows from the induction hypothesis. Multiplying out and canceling terms gives

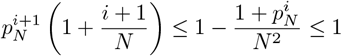

Taking 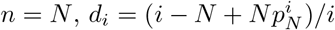 for *i* ∈ {2, 3, …, }, and applying Eq (A.1), we see

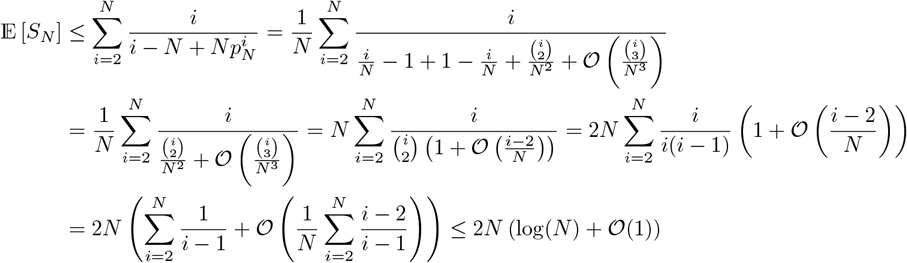

Dividing through by 2*N* log(*N*) and taking *N* → ∞ gives the theorem.

Let *B*^WF^(*n, N*_*e*_) denote the random total size of the tree of a sample of size *n* from a population evolving according to the Wright-Fisher model with effective size *N*_*e*_ and time measured in generations, so that *B*^WF^(*N*_*e*_, *N*_*e*_) is the random size of the tree of the whole population, and recall Thm A.2. Denote with *B*^WT^(*n, N*_*e*_) the total tree length for a sample of size *n* drawn from a population evolving according to the Wakeley-Takahashi model with effective size *N*_*e*_ and time measured in generations. We claim that, for any *x >* 0,

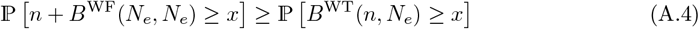

since *B*^WT^(*n, N*_*e*_) = *n*+*B*^WF^(*M, N*_*e*_) for some *M* ∈{1, 2, …, *N*_*e*_} where *M* is the random number of remaining lines after the first transition in the Wakeley-Takahashi model.

For *n* ∈ ℕ, *b >* 1 fixed, and some positive random variable *X*,

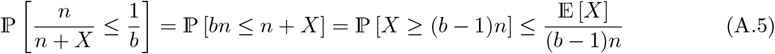

by the Markov inequality.

Let 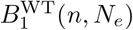 denote the random total length of external branches for a sample of size *n* drawn from a population evolving according to the Wakeley-Takahashi model with effective size *N*_*e*_ and time measured in generations. Note that 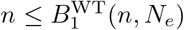. Taking *x* = *bn* for some fixed *b >* 1 then

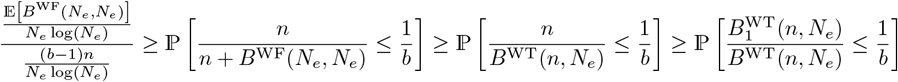

where the first inequality follows from Eq (A.5), the second inequality follows from Eq (A.4), and the third follows from 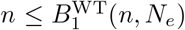. Then by Thm A.2 increasing the sample size *n*, so that *n/* (*N*_*e*_ log(*N*_*e*_)) → ∞ corresponding to our argument above, would tend to increase the relative length of external branches in the Wakeley-Takahashi model.

## B A lower bound on ∥Q_*N*_ − Q∥

The result we obtain in this section, a lower bound on the matrix norm ∥**Q**_*N*_ − **Q**∥ as defined in (Möhle, 2000) may be well known, but as we did not find a reference for it we include it here.

For ease of presentation we now present the notation of Möhle (2000) that we require. Throughout this subsection, the matrix norm ∥**A∥** for a matrix **A** = (*a*_*ij*_) is given by ∥**A∥** = sup_*i*_ ∑_*j*_ |*a*_*ij*_|. Let *ν*_*i*_ denote the random number of surviving offspring of the *i*th individual (arbitrarily labelled) in a haploid population of fixed size *N*, where the *ν*_1_, …, *ν*_*N*_ are exchangeable. Denote by **Q**_*N*_ the infinitesimal generator of the Kingman-coalescent, and by **Q**_*N*_ = (**P**_*N*_ − **I**)*/c*_*N*_, where **P**_*N*_ is the one-step transition matrix of the (pre-limiting) ancestral process, **I** is the identity matrix, and *c*_*N*_ as defined in Def 2.4. Let **Q** denote the infinitesimal generator of the Kingman-coalescent.

An upper bound on ∥**Q**_*N*_ − **Q**∥ is given in Cor 3 in Möhle (2000). Here we obtain a lower bound on ∥**Q**_*N*_ − **Q**∥ assuming a haploid Wright-Fisher population of fixed size *N* : referring to Möhle (2000) for notation, for *b*_1_ ≥ … ≥ *b*_*r*_ ≥ 1 and *b* = *b*_1_ + · · · + *b*_*r*_ (recall Eq (2.3))

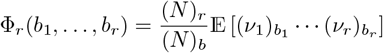

and from the proof of Lemma 2 in Möhle (2000), we have, with *G*(*a*) := 1 − Φ_*a*_(2, 1, …, 1)*/c*_*N*_,

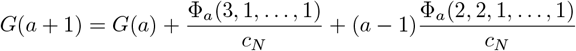

The factorial moments of a multinomial with equal probabilities *p*_*i*_ = 1*/N* for assigning an offspring to parent *i*,

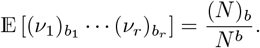

We see, for the Wright-Fisher model,

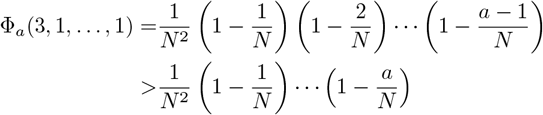

and restricting 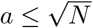 we see

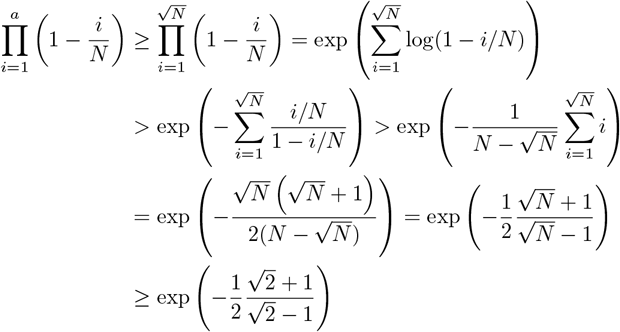

for *N* ≥ 2. Therefore, for 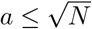,

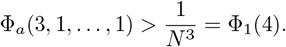

Considering Φ_*a*_(2, 2, 1, …, 1), the same calculation gives, for 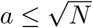,

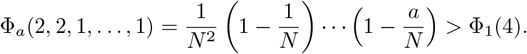

Then, from the proof of Lemma 2 in Möhle (2000), we have

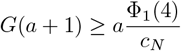

Applying Eq. 17 in Möhle (2000), i.e.

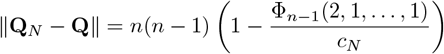

we conclude, that for the WF model with *c*_*N*_ = 1*/N*,

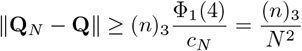

Then, by Example 1 of Möhle (2000), in the case of the Wright-Fisher model,

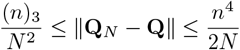

## C Comparing complete and incomplete Beta-coalescents

In this section we give a numerical example (see Figure 5) showing how estimates of mean relative branch lengths 𝔼 [*R*_*i*_(*n*)], recall (4.5), compare between the incomplete and the complete Beta-coalescent. The aim is to understand how inference based on the site-frequency spectrum might be influenced by the upper bound. The main result is that misspecifying the juvenile law (ignoring the upper bound) can lead to wrong conclusions. Recall the notation introduced in Section 4.

In Figure 5 we compare relative branch lengths sampled under different coalescents, by iteratively sampling trees under a given coalescent, and recording the relative branch lengths for each iteration, all for sample size *n* = 100. The coalescents considered are the Beta-coalescent of Schweinsberg (2003) (see Definition 2.2; red line), the Kingman coalescent (black line), and the incomplete Beta-coalescent (recall (3.12)) for values of *K* as shown. For all the Beta-coalescents *α* = 1.01. Figure 5 clearly shows that for small *K*, the site-frequency spectrum predicted by the incomplete Beta-coalescent resembles that of the Kingman coalescent more than that of the complete Beta-coalescent. This should not be surprising given the form of the coalescent rate for the incomplete Beta coalescent (recall (3.12)), where we integrate over the interval (0, *K/*(*K* + *m*_∞_)), so that smaller values of *K* translate to smaller probability of each line participating in a merger, reflecting the role of *K* in the prelimiting model (cf. Case 2 in Proposition 3.2). This conclusion holds for any *α*∈ (1, 2), and applying the complete Beta coalescent in inference, instead of the incomplete Beta coalescent, can lead to overestimating *α*. A ‘U-shaped’ (an excess of singletons and high-frequency variants relative to predictions of the Kingman-coalescent) is sometimes seen as a characteristic of Λ-coalescents (Birkner et al., 2013b; Freund et al., 2023). Our results show that, even though the offspring number distribution is highly skewed and in the domain of attraction of a multiple-merger coalescent, the predicted site-frequency spectrum is not necessarily U-shaped. Even though we have an explicit form of the cutoff point (*K/*(*K* + *m*_∞_)) of the incomplete Beta-coalescent, in inference one may view the stated coalescent as involving two parameters, *α* ∈ (1, 2), and (say) *γ* ∈ (0, 1], where *γ* represents the cutoff point. One could refer to the incomplete Beta-coalescent as the Beta(*γ*, 2 −*α, α*)-coalescent (and the Beta(2 −*α, α*)-coalescent is the Beta(*γ*, 2 −*α, α*)-coalescent when *γ* = 1). Given data one would then proceed to jointly estimate *α* and *γ*.

## D Comparing 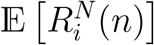 and 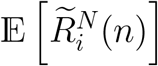

In this section we use simulations to investigate relative branch lengths read off trees sampled from a finite haploid panmictic population by conditioning on the population ancestry. This is done by simulating the evolution of a finite haploid population of fixed size *N* evolving according to Definition 2.3 and (4.1) and keeping track of the ancestry of each individual. Every now and then we take a random sample of size *n* and check if the sample coalesces, i.e. if a common ancestor of the sampled leaves exists (regardless of the ancestry of the individuals not sampled). If a (most recent) common ancestor of a given random sample is not found the sample is discarded and the population evolves further for some time until a new sample is drawn (independently of the previous one). If a common ancestor of a given sample is found, the ancestry of the sample is fixed, and the branch lengths are read off the fixed tree relating the sampled leaves. An estimate of the mean relative branch lengths is then obtained by repeating this a given number of times, each time starting from scratch with a new population. We denote the quantity thus approximated by 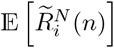. To our knowledge a coalescent that would approximate the trees sampled in this way from a large population has not been derived (and deriving such a coalescent is outside the scope of the present work). Therefore, we compare approximations of 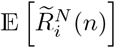 to approximations of 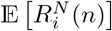 and the results can be found in Figure 6. Even though the approximations of 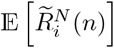 resp. 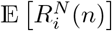 broadly agree there is a noticeable difference. Appendix F contains a brief description of an algorithm for approximating 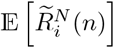, and Appendix E for approximating 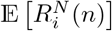.

## E Approximating 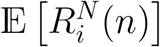

In this section we briefly describe an algorithm for approximating 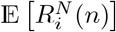 (recall (4.5) and Section 4).

We describe an algorithm for sampling branch lengths of a gene genealogy of a sample of size *n* from a finite haploid population of constant size *N*. Suppose the population evolves according to Definition 2.3. Our algorithm returns a realisation 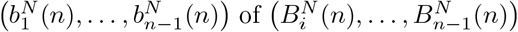 (recall (4.5) in Section 4). Since we are simulating a Markov sequence (a Markov process with a countable state space and evolving in discrete time) we only need to keep track of the current block sizes 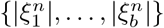 given that there are *b* blocks at the time where 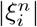 is the size (number of leaves the block is ancestral to) of block *i*, so that 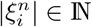 and 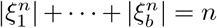. Let 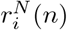 (initalised to zero) denote an estimate of 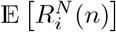 obtained in the following way:

1. Set each 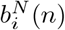 to zero
2. set each block size 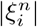 to one
3. while there are at least two blocks repeat the following steps in order
  a. update the branch lengths of the current blocks, i.e. 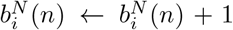 for 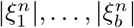 given that there are *b* blocks at the time
  b. sample a realisation *x*_1_, …, *x*_*N*_ of the juvenile numbers 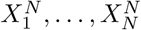 (using that *X*_*i*_ ~ ⌊*U*^−1*/α*^⌋ when the law of *X*_*i*_ is as in (4.4))
  c. sample number of blocks per family according to a multivariate hypergeometric with parameters *b* (the current number of blocks) and *x*_1_, …, *x*_*N*_
  d. merge blocks at random according to the numbers sampled in (c); for example suppose blocks of size 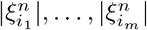 merge, then the continuing block has size 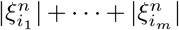
4. update the estimate 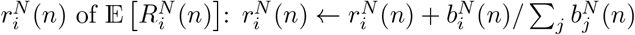
5. after repeating steps (1) to (4) a given number (𝕄) of times return 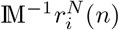 as an approximation of 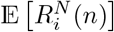

## F Approximating 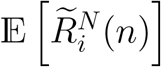

In this section we briefly describe an algorithm for approximating 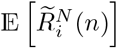 (see Figure 6 in Appendix D).

Let 𝔸^(*N,n*)^ ≡ (*A*_*i*_(*g*))_*g*∈ℕ ∪{0},*i*∈[*N*]_ denote the ancestry of the population. In our construction the population of *N* haploid individuals is partitioned into *N* levels. At any given time each level is occupied by one individual. Let *A*_*i*_(*g*) ∈ [*N*] be the level of the immediate ancestor of the individual occupying level *i* at time *g*. Set *A*_*i*_(0) = *i* for *i* ∈ [*N*]. If *A*_*i*_(*g*) = *A*_*j*_(*g*) for *i j* the individuals occupying levels *i* and *j* at time *g* derive from the same immediate ancestor. If the individual on level *i* at time *g* produces *k* surviving offspring then 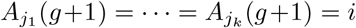. Each individual ‘points’ to its’ immediate ancestor (see Figure 7). A *complete* sample tree is one where the leaves have a common ancestor in the given population ancestry (e.g. the ‘blue’ sample in Figure 7). Let 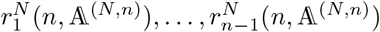 denote the realised relative branch lengths of a complete sample tree whose ancestry is given by 𝔸^(*N,n*)^. We approximate, for *i* ∈ {1, 2, …, *n* − 1},

**Figure 7:**
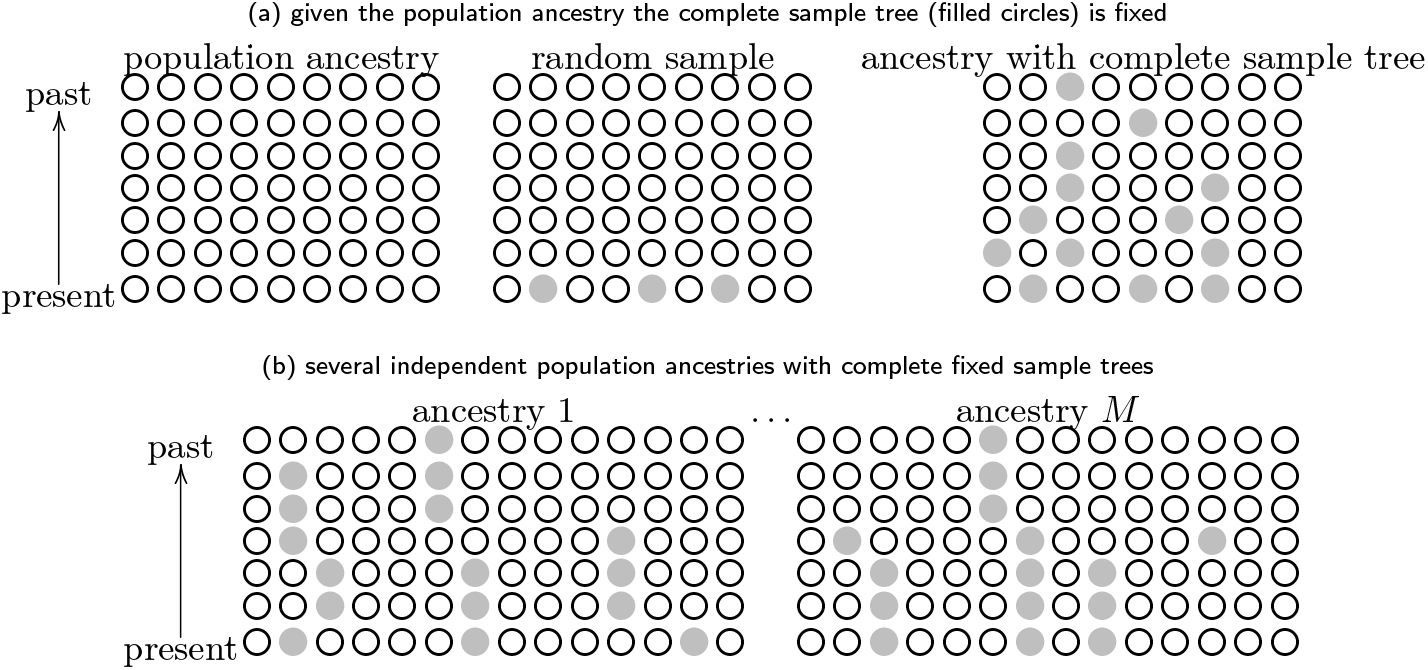
Illustrating the algorithm for generating conditional (quenched) gene genealogies. We take ‘ancestry’ to mean that we know the ancestral relations of the individuals in the population at any time. The filled circles 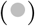 represent sampled gene copies (*n* = 3) or gene copies ancestral to the sampled ones. The empty circles (°) are gene co pies that are not ancestors of the sampled ones. Figure 6 holds examples of approximations of 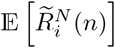 using the method illustrated here

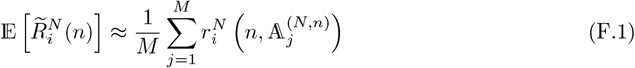

where *M* is the number of experiments, the number of realised ancestries 𝔸^(*N,n*)^.

We summarize the algorithm for approximating 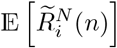 :

1. For each experiment:
  a. initialise the ancestry to *A*_*i*_(0) = *i* for *i* ∈ [*N*]
  b. until a complete sample tree is found:
    i. draw a random sample, i.e. sample *n* of *N* levels at the most recent time
    ii. check if the tree of the given sample is complete, if not discard the sample and record the ancestry of a new set of surviving offspring:
      A. sample numbers of potential offspring *X*_1_, …, *X*_*N*_
      B. given *X*_1_, …, *X*_*N*_ potential offspring sample the surviving offspring uniformly at random without replacement and update the ancestry; if the individual on level *i* at time *g* produces *k* surviving offspring then 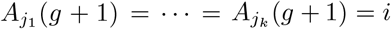.
  c. given a complete tree read the branch lengths off the tree and merge blocks according to the ancestry; suppose two blocks have ancestors on levels *i* and *j* at time *g*, if *A*_*i*_(*g*) = *A*_*j*_(*g*) the blocks are merged;
  d. given the branch lengths 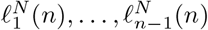 of a complete tree update the estimate 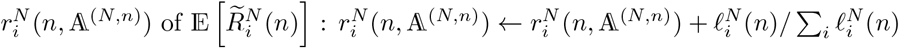
2. From *M* realised ancestries 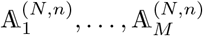 return (F.1) as an approximation of 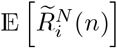 for *i* = 1, 2, …, *n* − 1

The idea is illustrated in Figure 7; (a) given a population ancestry the complete sample tree is fixed as soon as the identity of the sampled gene copies is known; (b) for each population ancestry we read the branch lengths off one sample tree, and then average the branch lengths over population ancestries.

